# An engineered monomer binding-protein for *α*-synuclein efficiently inhibits the proliferation of amyloid fibrils

**DOI:** 10.1101/568501

**Authors:** Emil D. Agerschou, Theodora Saridaki, Patrick Flagmeier, Céline Galvagnion, Daniel Komnig, Akansha Nagpal, Natalie Gasterich, Laetitia Heid, Vibha Prasad, Hamed Shaykhalishahi, Aaron Voigt, Dieter Willbold, Christopher M. Dobson, Björn H. Falkenburger, Wolfgang Hoyer, Alexander K. Buell

## Abstract

Removing or preventing the formation of *α*-synuclein aggregates is a plausible strategy against Parkinson’s disease. To this end we have engineered the *β*-wrapin AS69 to bind monomeric *α*-synuclein with high affinity. In cultured cells, AS69 reduced the occurrence of *α*-synuclein oligomers and of visible *α*-synuclein aggregates. In flies, AS69 reduced *α*-synuclein aggregates and the locomotor deficit resulting from *α*-synuclein expression in neuronal cells. In a mouse model based on the intracerebral injection of pre-formed *α*-synuclein seed fibrills (PFFs), AS69 co-injection reduced the density of dystrophic neurites observed three months later. In biophysical experiments in *vitro*, AS69 highly sub-stoichiometrically inhibited auto-catalytic secondary nucleation processes, even in the presence of a large excess of monomer. We present evidence that the AS69-*α*-synuclein complex, rather than the free AS69, is the inhibitory species responsible for sub-stoichiometric inhibition. These results represent a new paradigm that high affinity monomer binders can be strongly sub-stoichiometric inhibitors of nucleation processes.

## Introduction

Cytoplasmic aggregates of the protein *α*-synuclein are the pathological hallmark of Parkinson’s disease (PD) and other synucleinopathies ^1^. Point mutations in the *α*-synuclein gene or triplication of the *α*-synuclein locus are associated with famillial forms of PD, and the *α*-synuclein locus is a genetic risk factor for sporadic PD. Targeting *α*-synuclein pathology is therefore a plausible strategy to stop disease progression in PD. Since *α*-synuclein aggregate pathology was demonstrated to propagate from neuron to neuron ^2^, recent work has focused on understanding the cellular and molecular events in this process. From a therapeutic perspective, *α*-synuclein aggregation is thought to be the underlying cause of PD and remains the focus of causal therapeutic strategies. The link between *α*-synuclein aggregation and PD has been known for two decades ^1,3^; however, the translation of this scientific discovery into a therapy has proven challenging. Since the first description of small molecules that inhibit *α*-synuclein aggregation ^4^, the search for promising compounds continues ^5–9^. While the first small molecules also inhibited the aggregation of tau and amyloid-β, more recent compounds bind *α*-synuclein more selectively and showed reduced *α*-synuclein toxicity in mouse models of PD ^7^.

We have taken a different strategy by engineering a protein, the *β*-wrapin AS69, to induce the formation of a *β*-hairpin in monomeric *α*-synuclein upon binding ^10^. AS69, which was selected by phage display ^10^ from protein libraries based on ZAβ3, an affibody against the amyloid-β peptide ^11–13^, thus not only binds *α*-synuclein with high affinity, but induces a specific conformational change - akin to molecular chaperones ^14^. AS69 induces local folding of the region comprising residues 37-54 into a *β*-hairpin conformation in the otherwise intrinsically disordered, monomeric *α*-synuclein, and inhibits the amyloid fibril formation of *α*-synuclein under conditions of vigorous shaking of the solution even at highly substoichiometric ratios ^10^. Amyloid fibril formation, however, is not a one-step process but can be decomposed into different individual steps, including primary and secondary nucleation and fibril elongation. With vigorous shaking, for instance, primary nucleation can occur readily at the air-water interface ^15^ and fibril fragmentation induced by the shaking amplifies the number of growth-competent fibril ends ^16^. In order to validate AS69 as a potential therapeutic agent we therefore tested its biological effects in cellular and animal models, and we found it to be a highly efficient inhibitor of *α*-synuclein aggregation and associated toxicity. In addition, we designed a set of experimental conditions to measure selectively the effect of AS69 on specific steps of *α*-synuclein aggregation. We found that AS69 is able to efficiently interfere with the auto-catalytic secondary nucleation process of *α*-synuclein amyloid fibril formation. This effect on secondary nucleation is observed even in the presence of a large excess of *α*-synuclein monomer, which is expected to sequester AS69 into inhibitor-monomer complexes. We show evidence that the secondary nucleation of *α*-synuclein can be inhibited by the *α*-synuclein-AS69 complex and that therefore the inhibitory effect of AS69 is unaffected by even large excess concentrations of free *α*-synuclein monomer.

## Results

### Coexpression of AS69 reduces the formation of *α*-synuclein oligomers in cell culture

First, we explored the effect of the expression of AS69 on the viability of living cells and the aggregation of *α*-synuclein in a cellular environment. In these model systems we not only expressed WT *α*-synuclein but also the A53T variant, which has been associated with familial PD and which produces aggregates more quickly than the WT protein ^3,17^. We first used bimolecular fluorescence complementation (BiFC) to detect *α*-synuclein oligomers in living HEK293T cells ^18^. Constructs of WT and A53T *α*-synuclein were tagged with the C-terminal segment of the fluorescent protein Venus (synuclein-VC) or with the complementary N-terminal segment of this protein (VN-synuclein) (Figure 1 a). Neither of the two Venus fragments shows significant fluorescence by itself, but together they can generate a functional fluorescent protein ^19^ and hence function as a reporter for aggregation. We then transfected HEK293T cells with both synuclein-VC and VN-synuclein, in addition to AS69 (or LacZ as a control) and determined by flow cytometry the fraction of cells that displayed Venus fluorescence (Figure 1 b). In the absence of AS69, the fraction of fluorescent cells was larger with the expression of A53T-*α*-synuclein than WT-*α*-synuclein (Figure 1 b, p<0.05, two-way ANOVA). Coexpression of AS69 with both variants reduced the fraction of fluorescent cells, and hence of *α*-synuclein aggregates, to similar levels (Figure 1 b, p<0.05 for WT and p<0.01 for A53T, two-way ANOVA). AS69 did not, however, reduce the total quantity of *α*-synuclein in the cells, as determined from immunoblots (Figure 1 c, note that only the upper band reports *α*-synuclein ^20^). This finding is consistent with the hypothesis that the effects of AS69 in this cellular model system result from the inhibition of aggregation, and not from an enhanced clearance of *α*-synuclein. We then probed the effects of AS69 on larger aggregates of *α*-synuclein by transfecting HEK293T cells with A53T-*α*-synuclein tagged with enhanced green fluorescent protein (EGFP) as previously described ^20–22^ (Figure 1 d). The distribution of EGFP within transfected cells was classified as homogenous, containing particles or unhealthy (rounded cells that in time-lapse microscopy were observed to subsequently undergo apoptosis). Co-expression of AS69 with A53T *α*-synuclein led to an increase in the fraction of cells with a homogenous distribution of EGFP and fewer cells showed *α*-synuclein particles relative to those cells without AS69 (Figure 1 d). These findings indicate that AS69 reduces *α*-synuclein oligomers and visible aggregates in cultured human cells.

**Figure 1:**
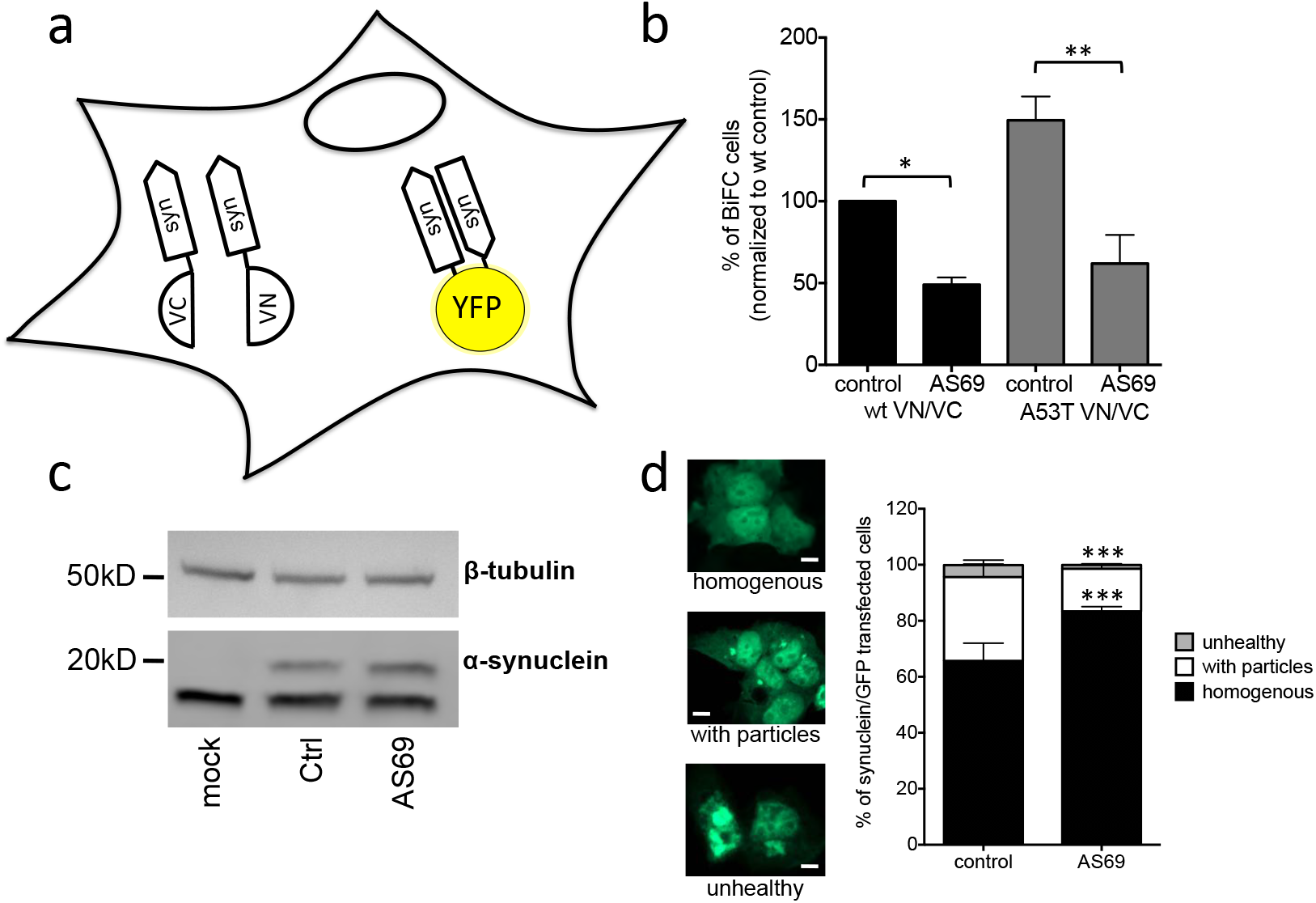
AS69 reduces the aggregation of *α*-synuclein in cellular models. **(a)** Schematic representation of bimolecular fluorescence complementation where *α*-synuclein is tagged by either the C-terminal (VC) or the N-terminal (VN) fragment of the Venus protein. In dimers or larger oligomers of *α*-synuclein, the two Venus fragments can form a functional fluorescent protein. **(b)** The percentage of fluorescent cells as determined by flow cytometry. HEK293T cells were transfected with *α*-synuclein (WT or A53T), fused to the VN or VC fragment and either LacZ (control) or AS69. Displayed are mean +/− SEM of n=3 independent experiments with n=100,000 cells analyzed per group in each experiment. **(c)** Immunoblots of lysates transfected with EGFP-tagged *α*-synuclein and in addition AS69 or LacZ (control), developed with antibodies against *α*-synuclein (bottom) and *β*-tubulin (top), the latter as a loading control. Similar findings were obtained in n=3 independent blots, where quantification showed no significant difference between Ctrl. and AS69 (t-test). **(d)** HEK293T cells were transfected with EGFP-tagged *α*-synuclein and the distribution of fluorescence was determined; the data shown are mean +/− SEM of n=3 independent experiments with n=300 cells classified per group in each experiment.

### Coexpression of AS69 rescues A53T *α*-synuclein dependent phenotype in Drosophila melanogaster

Subsequently we tested the effects that AS69 has in Drosophila melanogaster (fruit flies) expressing A53T-*α*-synuclein in neurons (Figure 2). In the absence of AS69, these flies show a progressive reduction in the spontaneous climbing (*i.e*. neuronal impairment) between 15 and 25 days of age ^20, 23^. We then generated flies co-expressing either AS69 or GFP (as a control) with A53T *α*-synuclein in neurons. Flies expressing AS69 and A53T *α*-synuclein showed preserved climbing behaviour (Figure 2 b, two-way ANOVA), demonstrating that neuronal expression of AS69 reduces the phenotype in this fly model of A53T *α*-synuclein toxicity. In order to determine whether or not the observed effect of AS69 on climbing behaviour could result from a reduction in the number of *α*-synuclein aggregates, we used flies expressing in all neurons one copy of A53T-*α*-synuclein fused to VC, one copy of A53T-*α*-synuclein fused VN ^24^, and in addition AS69 or “always early RNAi” as a control. Aggregates of *α*-synuclein were quantified by a filter trap assay in which urea treated lysates of fly heads were passed through a membrane and the quantity of *α*-synuclein aggregates retained in the membrane was detected by antibodies raised against *α*-synuclein (Figure 2 c). We found that the quantity of aggregates retained in the filter was much smaller in lysates from flies coexpressing AS69 and A53T-*α*-synuclein than in lysates from flies only expressing A53T-*α*-synuclein (Figure 2 d). This finding confirms that AS69 reduces high molecular weight aggregates of *α*-synuclein in neuronal cells of Drosophila melanogaster.

**Figure 2:**
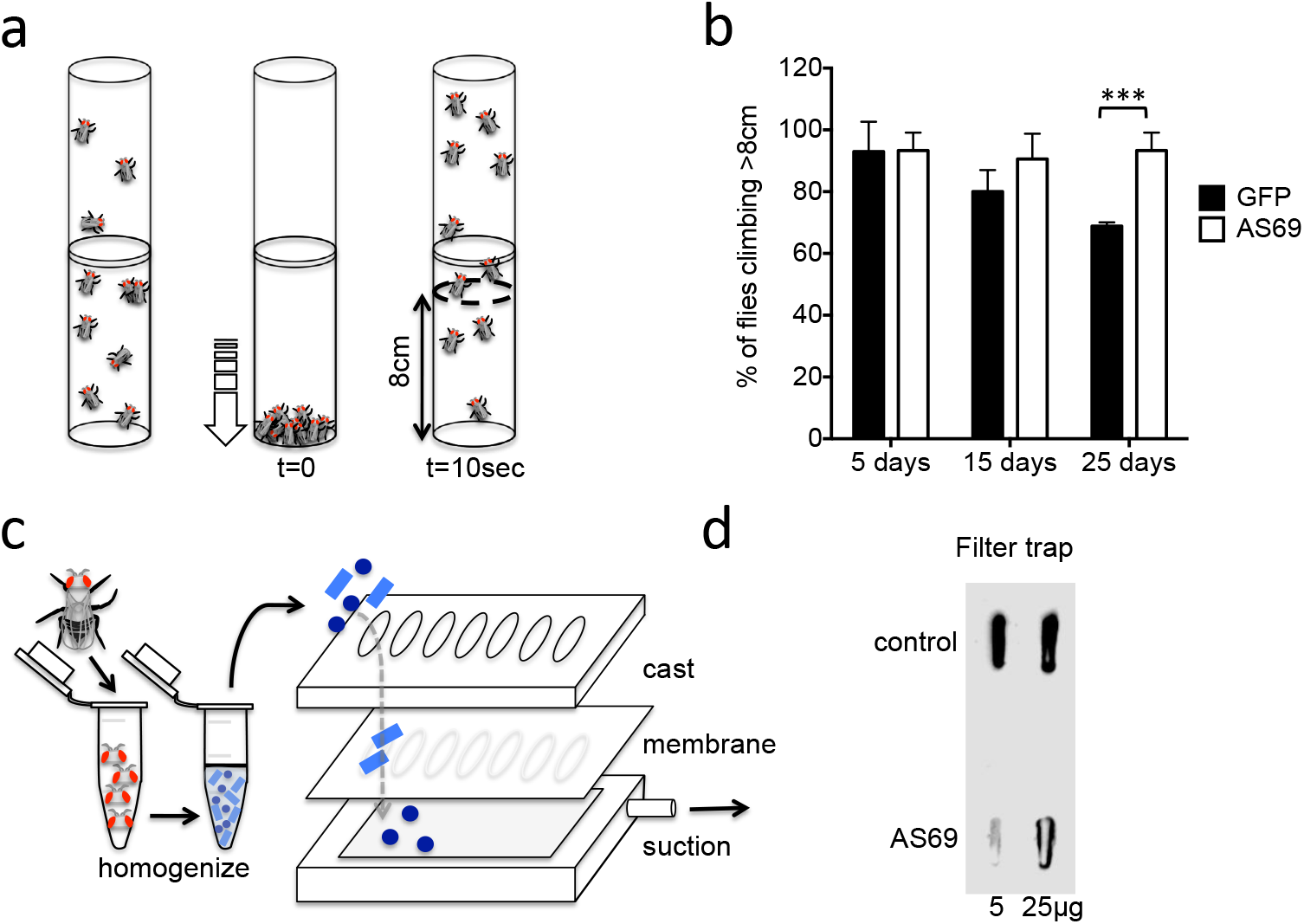
AS69 rescues the motor phenotype and reduces *α*-synuclein aggregation in *Drosophila melanogaster*. **(a)** Schematic representation of the climbing assay. The vials are tapped to move the flies to the base of the vial, and thereafter the flies climb towards the top of the vial; in this experiment the number of flies climbing 8 cm in 10 s was determined. **(b)** Performance in the climbing assay of Drosophila melanogaster expressing A53T-*α*-synuclein and either AS69 or GFP in neurons. At each time point, n = 30 flies were assayed per genotype; similar findings were observed for 8 different lines expressing AS69. **(c)** Schematic representation of the filter trap assay in which aggregates in the protein lysate are retained by a membrane, which is subsequently developed in the same manner as an immunoblot. **(d)** Results of the filter trap assay from lysates of control flies and flies expressing AS69 in addition to A53T-*α*-synuclein in all neurons. Two different quantities of the protein lysate were applied in each case, 5 and 25 *μ*g; similar findings were observed in n = 3 independent experiments.

### AS69 reduces Lewy neurites in a mouse model of PD

In order to investigate if the protective effects of AS69 could be extended to vertebrates, we tested the effects of AS69 in a mouse model based on the stereotactic injection of *α*-synuclein pre-formed seed fibrils (PFFs) into the brain of transgenic mice expressing A30P-*α*-synuclein under control of the neuronal Thy1 promoter. This model is similar to previous models using transgenic mice expressing WT and A53T-*α*-synuclein ^25–27^. Mice were analysed 3 months after PFFs were injected into the cortex and striatum using a single trajectory (Figure 3 a). In striatal sections stained for phosphorylated *α*-synuclein we observed large inclusion bodies in neuronal somata reminiscent of Lewy bodies (Figure 3 b, arrow heads), and dystrophic neurites loaded with phospho-synuclein positive material reminiscent of Lewy neurites (Figure 3 b, arrows). Since the Lewy neurite pathology (LN) was more frequent and more easily discriminated from background staining we quantified the density of LN in striatal sections 180 *μ*m rostral to the anterior commissure using stereological methods. Some LN were detected in animals injected with vehicle only (Figure 3 c), which was expected given that they were A30P *α*-synuclein transgenic mice. The density of LN was greatly increased by PFF injection (Figure 3 c), indicating that PFF can seed *α*-synuclein aggregates also in A30P-*α*-synuclein transgenic mice. The density of LN was significantly smaller in mice where AS69 was co-injected with the PFF (Figure 3 c), confirming that AS69 can reduce *α*-synuclein pathology in mice.

**Figure 3:**
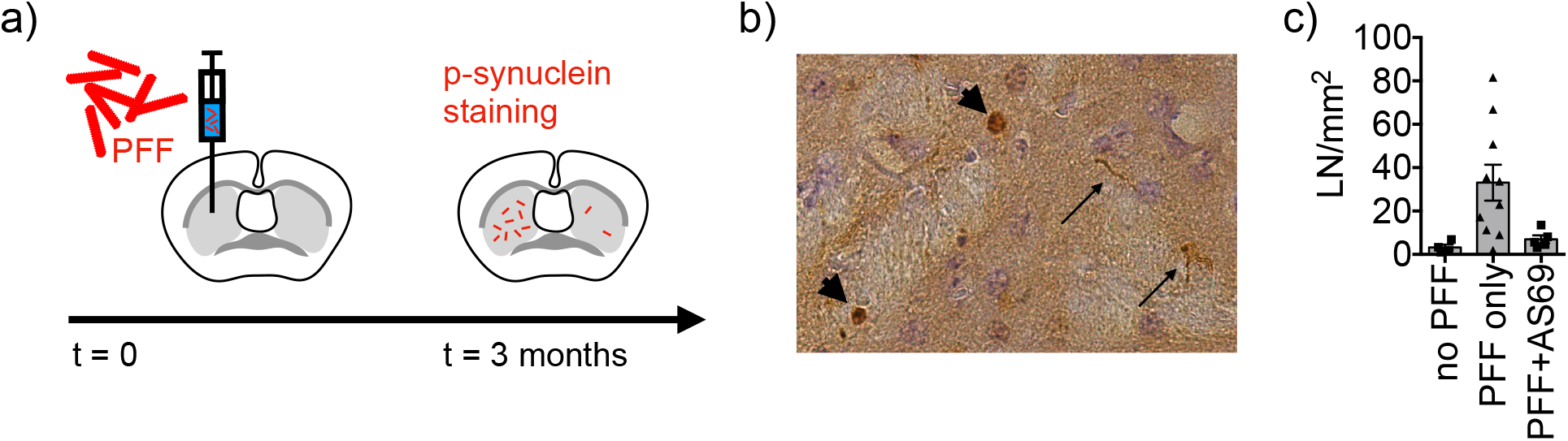
AS69 reduces synuclein pathology in a mouse model. **(a)** Experimental paradigm: Preformed *α*-synuclein fibrils (PFF), vehicle control, or PFF with AS69 were stereotactically injected into the left striatum of 12-week old A30P-*α*-synuclein transgenic mice. Mice were analysed 3 months later by staining striatal slices for phosphorylated *α*-synuclein. Dystrophic, *α*-synuclein-positive neurites were termed Lewy neurites and quantified in the striatal slice comprising the anterior commissure. **(b)** Image of striatal slice stained against phosphorylated *α*-synuclein. Neurons with strong staining indicated by arrowhead. Dystrophic neurites with positive staining indicated by long arrows. **(c)** Quantification of dystrophic neurites (LN) in the ipsilateral striatum from n = 4 animals for vehicle control (no PFF), n = 10 animals for PFF only, and n = 5 animals for PFF+AS69. One animal in the PFF+AS69 group was excluded because it displayed an abnormally high density of aggregates both on the injected and on the non-injected side (140 LN/mm^2^). Markers represent values for each animal. Bars represent mean +/− SEM. One-way ANOVA was significant (p = 0.0250), Holm-Sidaks multiple comparisons test revealed significant differences between “no PFF” and “PFF only” and between “PFF only” and “PFF+AS69”.

### AS69 stoichiometrically inhibits the elongation of *α*-synuclein fibrils

We next set out to elucidate the origin of the remarkable ability of AS69 to inhibit *α*-synuclein aggregate formation in cells and in *vivo* (Figures 1, 2 and 3) and amyloid fibril formation *in vitro* ^10^. To this end, we performed a detailed mechanistic analysis, where we examined the effect of AS69 on the growth and autocatalytic amplification of *α*-synuclein amyloid fibrils ^17, 28^. We first carried out experiments in the presence of micromolar concentrations (in monomer equivalents) of pre-formed seed fibrils of *α*-synuclein at neutral pH under quiescent conditions (Figure 4 a,b). We have shown previously that under these conditions only fibril elongation through the addition of monomeric *α*-synuclein to fibril ends occurs at detectable rates ^28^ and that the rate of *de novo* formation of fibrils is negligible. We therefore examined the effects of AS69 on fibril elongation and analyzed these data by fitting linear functions to the early stages of the aggregation time courses (see supplementary figure 1 and supplementary section 2 for details of the analysis). The results indicate that fibril elongation is indeed inhibited by AS69 in a stoichiometric concentration-dependent manner (Figure 4 c). In this experiment, both the seed fibrils and the AS69 compete for the monomeric *α*-synuclein and the relative affinities determine the kinetics and thermodynamics of the system.

**Figure 4:**
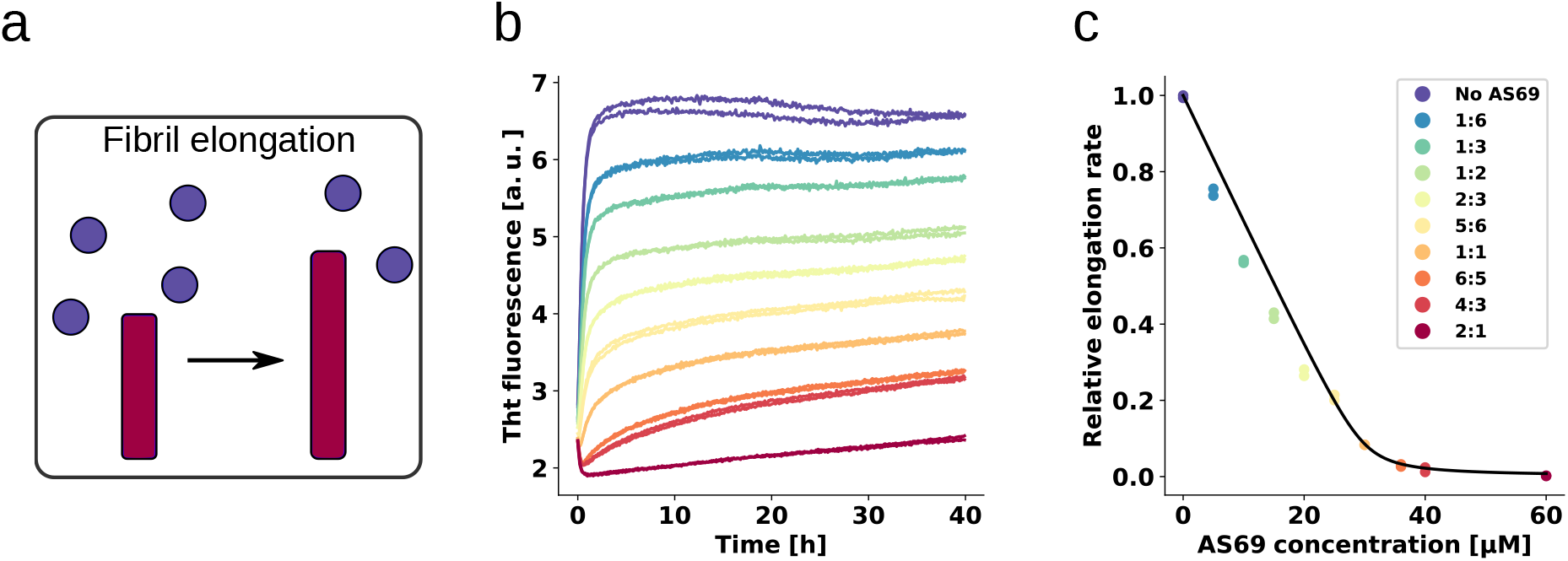
AS69 inhibits *α*-synuclein fibril elongation **(a)** Schematic representations of fibril elongation. **(b)** Change in ThT fluorescence when a 30 *μ*M solution of monomeric *α*-synuclein was incubated in the presence of 5 *μ*M pre-formed fibrils under quiescent conditions with increasing concentrations of AS69. **(c)** Relative rates of fibril elongation with increasing concentrations of AS69. The solid line corresponds to a simulation based on the affinity of AS69 for monomeric *α*-synuclein (240 nM^10^).

In order to obtain an estimate of the affinity of monomeric *α*-synuclein for the ends of fibrils, we performed elongation experiments at low monomer concentrations in the absence of AS69. We found evidence that the fibrils are able to elongate in the presence of 0.5 *μ*M monomeric *α*-synuclein (see supplementary figure 2), providing an upper bound of the critical concentration (which is formally equivalent to a dissociation constant, see supplementary section 2). Despite the similar affinity of monomeric *α*-synuclein for both fibril ends and AS69, the time scales of the two types of interactions are very different; monomeric *α*-synuclein was found to interact on a timescale of seconds with AS69, as seen by isothermal titration calorimetry (ITC) experiments ^10^, but to incorporate on a timescale of minutes to hours into free fibril ends (see Figure 4 b and ^28^). The slow kinetics of the latter process is partly due to the fact that the number of fibril ends is much smaller than the number of monomers ^28^, such that each fibril sequentially recruits many *α*-synuclein molecules. Therefore, the equilibrium between AS69 and *α*-synuclein should be rapidly established and perturbed only very slowly by the presence of the fibrils.

### The inhibition of fibril elongation is due to monomer sequestration

The initial fibril elongation rate as a function of AS69 concentration was found to follow closely the concentration of unbound *α*-synuclein across the entire range of concentrations of AS69 used in this study, as shown in Figure 4 c, where the prediction based on this simplifying assumption is shown as a solid line. The inhibition of fibril elongation can therefore be explained quantitatively by the sequestration of monomeric *α*-synuclein by AS69 and the assumption that the AS69:*α*-synuclein complex cannot be incorporated into the growing fibril. This conclusion is supported by the finding that the fibrils formed in the presence of increasing concentrations of AS69 are morphologically indistinguishable from the fibrils formed in the absence of AS69 (as judged from AFM images, see supplementary Figure 3). Our kinetic analysis of fibril elongation in the presence of AS69 does not, however, suggest a preferential interaction with fibril ends, as such an interaction can be expected to lead to a sub-stoichiometric inhibition of fibril elongation, which is not observed in our experiments. Indeed, the finding that the effect on elongation can be quantitatively described by considering only the interaction of AS69 with monomeric *α*-synuclein suggests a weak, if any, interaction of AS69 with fibrils. Furthermore, density gradient centrifugation (DGC) of samples containing only seeds and AS69 (see supplementary figure 4 a and b) did not show AS69 to co-migrate with large species to any significant extent under conditions that favour elongation. We also examined the effect that prolonged incubation of pre-formed seed fibrils with AS69 has on the seeding efficiency of the former (see supplementary figure 5) in order to probe whether the effect of AS69 in our mouse model could be explained by a strongly decreased seeding potential of the injected PFF due to their co-injection with AS69. We found that the seeding efficiency of untreated PFF, and of PFF that had been incubated for 24 h with a stoichiometric concentration of AS69, was very similar, indicating that co-incubation of seed fibrils with AS69 has no significant effect on their ability to recruit soluble *α*-synuclein.

### AS69 sub-stoichiometrically inhibits the amplification of *α*-synuclein fibrils

These findings clearly demonstrate that AS69 inhibits fibril elongation in a stoichiometric manner through monomer sequestration. Consequently, inhibition of fibril elongation cannot explain the previously observed substoichiometric inhibition of *α*-synuclein fibril formation by AS69 ^10^. We therefore performed seeded experiments under mildly acidic solution conditions in the presence of very low concentrations of pre-formed fibrils (nM monomer equivalents) under quiescent conditions (Figure 5 a, b) ^28, 29^. Under those solution conditions, seeded aggregation has been shown to consist of two processes in addition to fibril elongation, namely secondary nucleation, which increases the number of growth competent fibril ends, and higher order assembly (“flocculation”), which decreases the overall aggregation rate by reducing the number of accessible fibrils through their burial within higher order aggregates ^28^. We find that under these solution conditions, where only growth and secondary nucleation contribute to the increase in fibril mass and number, respectively, the seeded aggregation is inhibited in a strongly sub-stoichiometric manner (Figure 5 b,c). We analysed these data to determine the maximum rate of aggregation (see supplementary figure 6 and supplementary section 2 for details) using the framework from ^30^. Based on recent results on the concentration-dependence of autocatalytic secondary nucleation of *α*-synuclein amyloid fibrils ^29^, we have calculated the predicted inhibitory effect due to monomer sequestration by AS69 in Figure 5 c). We find that, unlike the case of fibril elongation, monomer sequestration cannot explain the extent of inhibition, even by assuming a very high reaction order of 5 (i.e. a dependence of the rate of secondary nucleation on the 5th power of the free monomer concentration; 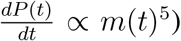 which is not compatible with recent results, showing that secondary nucleation depends only weakly on the concentration of free monomer ^29^. Even in this unlikely scenario, the very strong inhibitory effect of low AS69 concentrations cannot be explained by monomer depletion.

**Figure 5:**
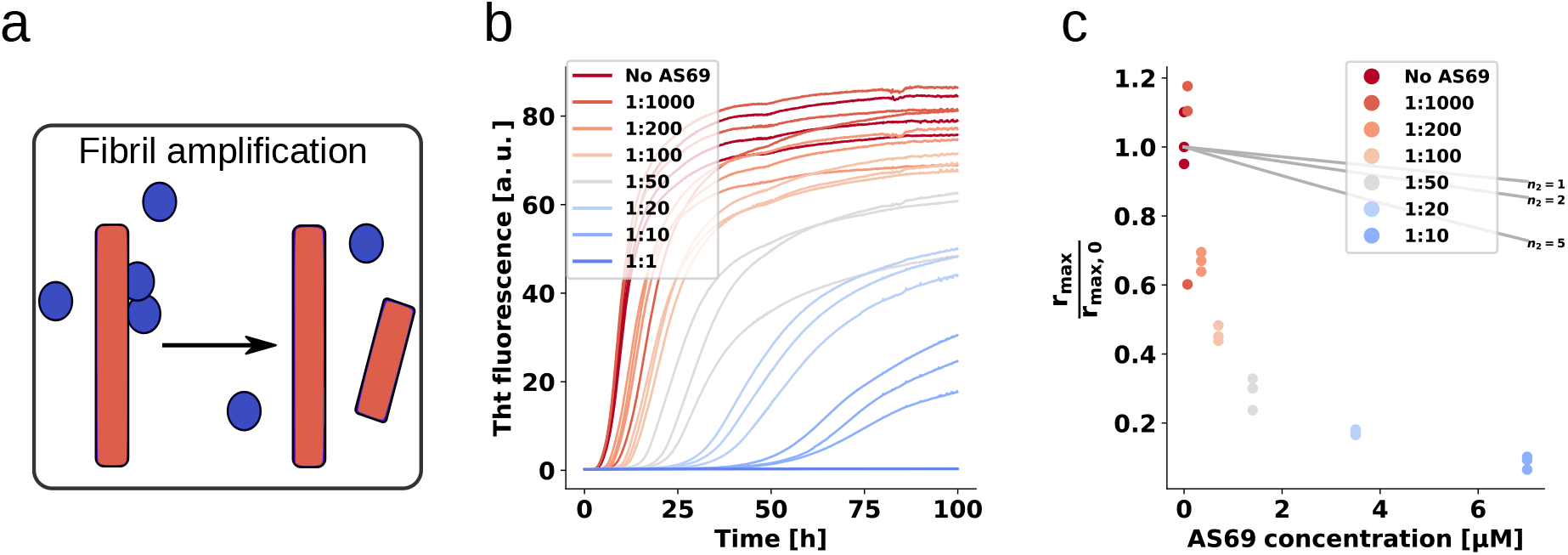
AS69 inhibits *α*-synuclein fibril amplification. **(a)** Schematic representation of fibril amplification. **(b)** Change in ThT fluorescence intensity when a 70 *μ*M solution of monomeric *α*-synuclein was incubated with increasing concentrations of AS69 in acetate buffer (pH 5.0) under quiescent conditions. **(c)** Relative rate of fibril amplification as a function of the concentration of AS69.

### The sub-stoichiometric inhibition of fibril amplification is not due to interaction with the fibril surface

We have previously been able to rationalise the inhibition of the secondary nucleation of *α*-synuclein by the homologous protein *β*-synuclein through a competition for binding sites on the surface of the fibrils ^31^. Here we find that AS69 is a significantly more efficient inhibitor of the autocatalytic amplification of *α*-synuclein amyloid fibrils than *β*-synuclein (a similar degree of inhibition is achieved with a ten fold lower concentration ratio). This result is particularly interesting in the light of the fact that AS69 binds efficiently to monomeric *α*-synuclein, whereas we found no evidence for a relevant direct interaction between the monomeric forms of *α*- and *β*-synuclein, given the complete absence of any inhibitory effect of *β*-synuclein on the elongation of *α*-synuclein fibrils ^31^. Therefore, despite the fact that the vast majority of the AS69 is bound within a complex with monomeric *α*-synuclein, AS69 is an efficient sub-stoichiometric inhibitor of the secondary nucleation of *α*-synuclein. This finding suggests that in addition to inhibiting through competition for nucleation sites on the fibril surface, AS69 or its complex with *α*-synuclein could interact directly with intermediates of the secondary nucleation process. To investigate whether AS69 binds to the fibril surface under these secondary nucleation-inducing solution conditions, we performed additional DGC experiments. Co-migration in the density gradient of AS69 with fibrils, which would imply direct interactions between these species, was undetectable (see Supplementary Figure 7 a,b and c). If AS69 were able to inhibit secondary nucleation through binding to the fibril surface in the presence of a large excess of monomer, its affinity to fibril surfaces would need to be much higher than to monomeric *α*-synuclein. This implies that under the conditions of the DGC experiments which were performed in the absence of monomeric *α*-synuclein, all binding sites on the fibrils should be occupied. Therefore, the absence of detectable binding implies either a weak affinity for fibrils or a very low stoichiometry, i.e. a very low density of binding sites for AS69 on the fibril surface.

### AS69 binds to stable *α*-synuclein oligomers with comparable affinity as to monomers

We next tested whether binding of AS69 to oligomeric states of *α*-synuclein could explain the efficient inhibition of secondary nucleation. The heterogeneous and often transient nature of oligomeric intermediates on the pathway to the formation of amyloid fibrils makes any interaction between such species and AS69 difficult to probe. However, monomeric *α*-synuclein can be converted into kinetically stable oligomers that can be studied in isolation, because they do not readily convert into amyloid fibrils ^32^. Despite the fact that these species are not likely to be fibril precursors, they are intermediate in size and structure between monomeric and fibrillar *α*-synuclein and hence can serve as a model for AS69 binding to *α*-synuclein oligomers. Using microscale thermophoresis (MST ^33^), we were able to confirm the binding of AS69 to both monomeric (see Supplementary Figure 8 a) and oligomeric *α*-synuclein (see Supplementary Figure 8 b) and provide estimates of the respective binding affinities (ca. 300 nM for monomeric and ca. 30 nM for oligomeric *α*-synuclein). The finding that AS69 is able to inhibit secondary nucleation in a highly sub-stoichiometric manner in the presence of a large excess of free monomer, to which it binds with high affinity, necessitates that the interactions of AS69 with aggregation intermediates must be of significantly higher affinity, if they are to explain the inhibition. Otherwise the monomer would out-compete the aggregation intermediate for AS69 binding, due to the much lower concentration of the latter. An estimate (see supplementary section 2 for details) suggests that the affinity of AS69 for aggregation intermediates would need to be several orders of magnitude higher than to *α*-synuclein monomer in order to explain an inhibitory effect of the observed magnitude. This required affinity is indeed much higher than the affinity we have determined here for an oligomeric state of *α*-synuclein.

### The covalent complex of AS69 and *α*-synuclein efficiently inhibits secondary nucleation

The analysis described in the previous section suggests, therefore, that the *α*-synuclein:AS69 complex itself could be the inhibitory species. The population of this complex is sufficiently high, even at low ratios of AS69:*α*-synuclein, to interact with a considerable fraction of aggregation intermediates. It is possible, therefore, that while the AS69:*α*-synuclein complex is unable to incorporate into a fibril end (see above), it can interact with oligomeric fibril precursors and block their conversion into fibrils.

We tested this hypothesis by producing a molecular construct whereby *α*-synuclein and AS69 are linked together with a flexible glycine tether that allows the formation of an intramolecular complex (AS69fusASN). The formation of the intramolecular complex was verified by performing CD spectroscopy at 222 nm over a the temperature range from 10 to 90°C and fitting the data to a two-state model ^34^ (see supplementary figure 9 and table). Both at neutral and mildly acidic pH, the fusion construct AS69fusASN has a higher thermal stability than the free AS69 and, indeed, as the stoichiometric mixture of AS69 and *α*-synuclein. The difference in melting temperatures between the covalent and non-covalent complex can be explained by the differences in the entropy of binding, which is more unfavourable in the case of the non-covalent complex, given the loss of three degrees of freedom of translational motion upon binding.

We performed weakly seeded aggregation experiments under conditions where secondary nucleation leads to the amplification of the added seed fibrils (see above) at different concentrations of AS69 (Figure 6 a) as well as AS69-*α*-syn complex (Figure 6 b). We found that the pre-formed complex is an equally efficient inhibitor as the free AS69 under secondary nucleation conditions (Figure 6 e). These results provide strong support of our hypothesis that the AS69-*α*-synuclein complex, covalent or non-covalent, is the species that is responsible for the sub-stoichiometric inhibition of secondary nucleation. Therefore, we propose a model whereby rather than requiring the binding of free AS69 to an aggregation intermediate, the AS69:*α*-synuclein complex is able to incorporate into a fibril precursor and efficiently prevent it from undergoing the structural rearrangement required to transform into a growth-competent amyloid fibril.

**Figure 6:**
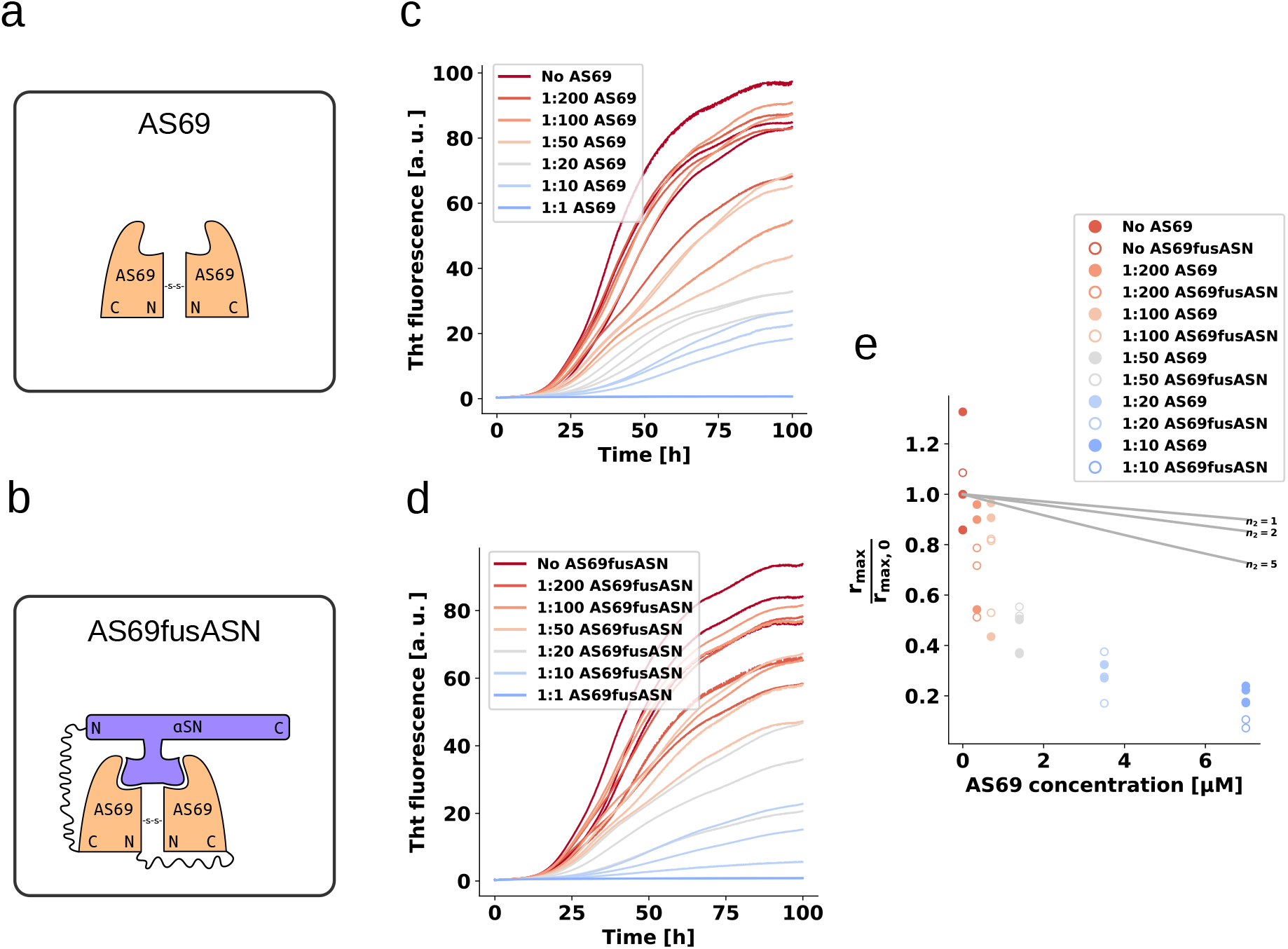
AS69 and AS69fusASN inhibits *α*-synuclein fibril amplification to similar extent. **(a)** and **(b)** Schematic representation of AS69 and AS69fusASN respectively. **(c)**, **(d)** Change in ThT fluorescence when a 70 *μ*M solution of monomeric *α*-synuclein was incubated with increasing concentrations of AS69 or AS69fusASN respectively in sodium acetate buffer (pH 5.0) under quiescent conditions. **(e)** Relative maximum rate of elongation as a function of the concentration of AS69 (closed circles) and AS69fusASN (open circles).

**Table 1:**
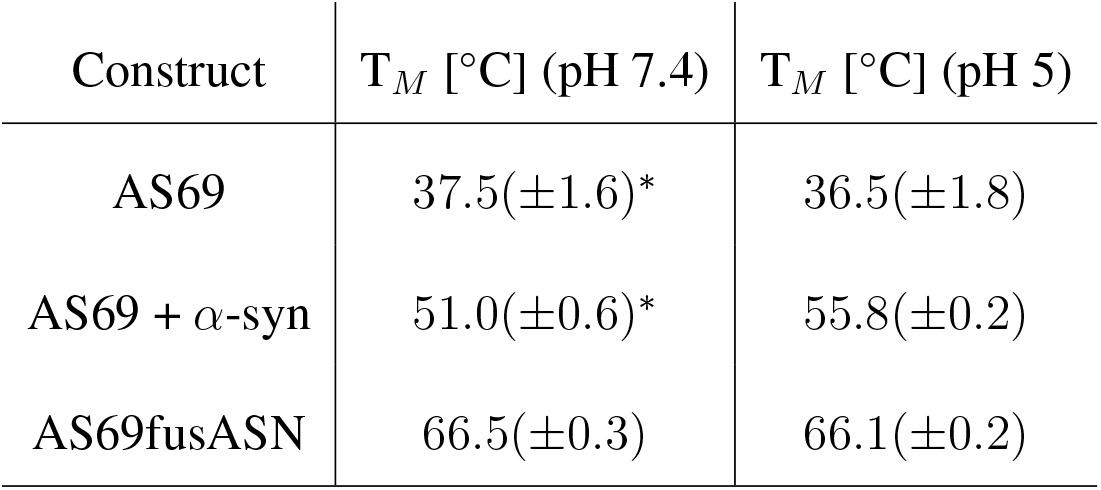
Selected parameters obtained from fitting the CD melting curves, the error in brackets represents the error obtained from fitting. * Data from ^35^ was refitted to obtain the numerical values.

| Construct | $T_M$ [°C] (pH 7.4) | $T_M$ [°C] (pH 5) |
| --- | --- | --- |
| AS69 | 37.5($\pm 1.6$)* | 36.5($\pm 1.8$) |
| AS69 + α-syn | 51.0($\pm 0.6$)* | 55.8($\pm 0.2$) |
| AS69fusASN | 66.5($\pm 0.3$) | 66.1($\pm 0.2$) |

## Discussion

The *β*-wrapin AS69 is a small engineered monomer binding protein that upon coupled folding-binding induces a local *β*-hairpin conformation in the region comprising amino acid residues 37-54 of otherwise intrinsically disordered monomeric *α*-synuclein. AS69 shows substoichiometric inhibition of *α*-synuclein aggregation in *vitro*, which is remarkable for a monomer binding-protein ^10^. Here, we show that potent aggregation inhibition of AS69 can be recapitulated in cell culture as well as animal models. In cell culture, AS69 reduced the amount of oligomers and visible aggregate particles (Figure 1). In flies, co-expression of AS69 led to reduced *α*-synuclein aggregation and rescue of the motor phenotype in the climbing assay (Figure 2). In a PD mouse model, the LN pathology induced by PFF injection was decreased when AS69 was co-injected (Figure 3). Our biophysical *in vitro* aggregation experiments under well-defined conditions enabled us to reveal two very distinct modes of inhibition of *α*-synuclein amyloid fibril formation by AS69. First, as expected for a monomer-binding species, AS69 inhibits fibril growth in a strictly stoichiometric manner, suggesting that the non-covalent AS69-*α*-synuclein complex is unable to add onto a fibril end and elongate the fibril. This is consistent with our results from density gradient centrifugation of the lack of a detectable interaction between AS69 and fibrils. Second, AS69 is found to be a very efficient inhibitor of secondary nucleation at highly sub-stoichiometric ratios. The overall result of our experimental and theoretical analysis is that this inhibitory effect is unlikely to stem from a direct interaction between the AS69 and either fibril surfaces or secondary nucleation intermediates. Such an interaction would need to be of an unrealistically higher affinity than the interaction between AS69 and *α*-synuclein monomer. A possible solution to this conundrum is presented by the hypothesis that indeed the AS69-*α*-synuclein complex is the inhibitory species. This hypothesis gains strong support from our finding that a covalently linked complex is an equally efficient inhibitor of secondary nucleation as the free AS69 molecule. It is important to note here that this proposed mode of action is very distinct from other types of inhibitory behavior reported previously. For example in the case of nanobodies raised against monomeric *α*-synuclein, at least stoichiometric amounts of the nanobodies are needed in order to interfere significantly with unseeded aggregation ^36^. In the case of molecular chaperones, on the other hand, sub-stoichiometric inhibitory behaviour has been reported previously ^37,38^, but it is usually found that these molecules do not interact significantly with the monomer, but rather bind specifically to aggregated states of the protein. Therefore, the AS69 affibody represents a new paradigm in the inhibition of amyloid fibril formation: strongly sub-stoichiometric inhibition by a tight monomer-binding species. In this scenario, not the inhibitor itself plays the role of a molecular chaperone, i.e. interaction with an on-pathway species and interfering with its further evolution, but rather the monomer-inhibitor complex acts as a chaperone. This mode of action represents a range of significant advantages over the other previously described modes of action (i.e. monomer sequestration and direct interaction with aggregation intermediates). First, it is rather straightforward to develop further molecules that bind to the monomeric forms of proteins, given that the latter are well-defined, reproducible and easy to handle. This simplicity is in contrast to the difficulty presented by targeting on-pathway aggregation intermediates which are difficult to isolate for the development of inhibitors. Second, binders of oligomeric aggregation intermediates can be expected to be less specific compared to binders of a well-defined monomeric state, as suggested by the existence of antibodies that interact with protofibrillar species independently of the protein from which they have formed ^39^. This lack of specificity can potentially lead to cross-reactivity and side effects. And third, the mode of inhibition here presented avoids the need for stoichiometric amounts of inhibitors that are usually required in the case of monomer sequestering species, resulting in a more efficient inhibition.

An inhibitor functioning according to this dual mode, i.e. being active both as free molecule and as a complex with monomeric *α*-synuclein, is expected to efficiently reduce *α*-synuclein aggregation in *vivo*. This is in agreement with the cell culture, fly, and mouse model data we present in this manuscript. The mode of action of AS69 in these three different biological models could be different. While AS69 is constantly expressed alongside *α*-synuclein in the cellular and fly models, only the initially co-injected AS69 is available for inhibition in the case of the mouse model. The relevant mode of inhibition in the cellular and fly model is therefore likely to be continued monomer sequestration, leading to the efficient suppression of the formation of toxic aggregates. On the other hand, the formation of LN pathology caused by cerebral injection of PFFs is likely to involve amplification of the injected PFFs by secondary processes, such as fibril fragmentation ^40^ or secondary nucleation ^28^. Based on the results from our detailed *in vitro* biophysical experiments, the strong reduction of *α*-synuclein pathology in the mouse model upon co-injection of AS69 could be explained by two possible mechanisms. First, the presence of the co-injected AS69 efficiently prevents the injected PFFs from growing and proliferating by binding to the endogenous soluble *α*-synuclein, according to the mechanisms deciphered in this work. The injected PFFs, deprived of their ability to grow and proliferate, would ultimately end up being degraded by the organism. Second, the high affinity of AS69 for soluble *α*-synuclein contributes to the dissolution and degradation of the injected PFFs, through a shift of the equilibrium between fibrils and free monomer induced by the sequestration of the free monomer. While we cannot rule out any of these two possibilities definitively, the latter process appears to be highly inefficient, given our finding that co-incubation of PFF with AS69 *in vitro* does not noticeably decrease the seeding capacity of the PFF. It is therefore likely that the effect of AS69 in our mouse model is also based on interference with fibril growth and amplification processes.

In conclusion, high affinity monomer binders displaying strong sub-stoichiometric inhibition of fibril formation represent attractive agents to interfere with pathological protein aggregation, due to their multiple inhibitory action.

## Acknowledgement

AKB thanks the Leverhulme Trust and the Parkinson’s and Movement Disorder Foundation (PMDF) for funding. PF thanks the Böhringer Ingelheim Fonds and the Studienstiftung des Deutschen Volkes for support. CG thanks the Alexander von Humboldt Foundation for support. This project has received funding from the European Research Council under the European Union’s Horizon 2020 research and innovation program, grant agreement No. 726368 (to WH). We thank Nadine Rösener for help with the density gradient centrifugation and Sabine Hamm for excellent technical assistance.

## References

1. Spillantini, M. G. et al. Alpha-synuclein in lewy bodies. Nature 388, 839–840 (1997).

2. Desplats, P. et al. Inclusion formation and neuronal cell death through neuron-to-neuron transmission of alpha-synuclein. Proc Natl Acad Sci U S A 106, 13010–13015 (2009).

3. Conway, K. A., Harper, J. D. & Lansbury, P. T. Accelerated in vitro fibril formation by a mutant alpha-synuclein linked to early-onset Parkinson disease. Nat Med 4, 1318–1320 (1998).

4. Masuda, M. et al. Small molecule inhibitors of α-synuclein filament assembly. Biochemistry 45, 6085–6094 (2006).

5. Wagner, J. et al. Anle138b: a novel oligomer modulator for disease-modifying therapy of neurodegenerative diseases such as prion and parkinson’s disease. Acta neuropathologica 125, 795–813 (2013).

6. Tóth, G. et al. Targeting the Intrinsically Disordered Structural Ensemble of α-Synuclein by Small Molecules as a Potential Therapeutic Strategy for Parkinson’s Disease. PloS one 9, e87133 (2014).

7. Wrasidlo, W. et al. A de novo compound targeting α-synuclein improves deficits in models of parkinson’s disease. Brain 139, 3217–3236 (2016).

8. Perni, M. et al. A natural product inhibits the initiation of α-synuclein aggregation and suppresses its toxicity. Proceedings of the National Academy of Sciences of the United States of America 114, E1009–E1017 (2017).

9. Kurnik, M. et al. Potent α-synuclein aggregation inhibitors, identified by high-throughput screening, mainly target the monomeric state. Cell chemical biology (2018).

10. Mirecka, E. A. et al. Sequestration of a β-hairpin for control of α-synuclein aggregation. Angew. Chem. Int. Ed. Engl. 53, 4227–4230 (2014).

11. Hoyer, W., Grönwall, C., Jonsson, A., Stahl, S. & Härd, T. Stabilization of a beta-hairpin in monomeric alzheimer’s amyloid-beta peptide inhibits amyloid formation. Proceedings of the National Academy of Sciences of the United States of America 105, 5099–5104 (2008).

12. Hoyer, W. & Härd, T. Interaction of alzheimer’s a beta peptide with an engineered binding protein–thermodynamics and kinetics of coupled folding-binding. Journal of molecular biology 378, 398–411 (2008).

13. Luheshi, L. M. et al. Sequestration of the abeta peptide prevents toxicity and promotes degradation in vivo. PLoS biology 8, e1000334 (2010).

14. Muchowski, P. J. & Wacker, J. L. Modulation of neurodegeneration by molecular chaperones. Nature reviews Neuroscience 6, 11–22 (2005).

15. Campioni, S. et al. The presence of an air-water interface affects formation and elongation of α-Synuclein fibrils. J. Am. Chem. Soc. 136, 2866–2875 (2014).

16. Xue, W.-F. et al. Fibril fragmentation enhances amyloid cytotoxicity. J Biol Chem 284, 34272–34282 (2009).

17. Flagmeier, P. et al. Mutations associated with familial parkinson’s disease alter the initiation and amplification steps of α-synuclein aggregation. Proceedings of the National Academy of Sciences of the United States of America 113, 10328–10333 (2016).

18. Falkenburger, B. H., Saridaki, T. & Dinter, E. Cellular models for parkinson’s disease. Journal of neurochemistry 139 Suppl 1, 121–130 (2016).

19. Bae, E.-J. et al. Glucocerebrosidase depletion enhances cell-to-cell transmission of α-synuclein. Nature communications 5, 4755 (2014).

20. Dinter, E. et al. Rab7 induces clearance of α-synuclein aggregates. Journal of neurochemistry 138, 758–774 (2016).

21. Opazo, F., Krenz, A., Heermann, S., Schulz, J. B. & Falkenburger, B. H. Accumulation and clearance of α-synuclein aggregates demonstrated by time-lapse imaging. Journal of neurochemistry 106, 529–540 (2008).

22. Karpinar, D. P. et al. Pre-fibrillar alpha-synuclein variants with impaired beta-structure increase neurotoxicity in Parkinson’s disease models. The EMBO Journal 28, 3256–3268 (2009).

23. Butler, E. K. et al. The Mitochondrial Chaperone Protein TRAP1 Mitigates α-Synuclein Toxicity. PLoS genetics 8, e1002488 (2012).

24. Prasad, V. et al. Monitoring α-synuclein multimerization in vivo. FASEB journal: official publication of the Federation of American Societies for Experimental Biology fj201800148RRR (2018).

25. Luk, K. C. et al. Pathological α-synuclein transmission initiates parkinson-like neurodegeneration in nontransgenic mice. Science (New York, N.Y.) 338, 949–953 (2012).

26. Sacino, A. N. et al. Induction of cns α-synuclein pathology by fibrillar and non-amyloidogenic recombinant α-synuclein. Acta neuropathologica communications 1, 38 (2013). Original DateCompleted: 20131122.

27. Masuda, H. et al. Antemortem detection of colonic [alpha]-synuclein pathology in a patient with pure autonomic failure. Journal of neurology 261, 2451 (2014).

28. Buell, A. K. et al. Solution conditions determine the relative importance of nucleation and growth processes in α-synuclein aggregation. Proc. Natl. Acad. Sci. U.S.A. 111, 7671–7676 (2014).

29. Gaspar, R. et al. Secondary nucleation of monomers on fibril surface dominates α-synuclein aggregation and provides autocatalytic amyloid amplification. Quarterly Reviews of Biophysics 50 (2017).

30. Cohen, S. I. A. et al. Nucleated polymerization with secondary pathways. I. time evolution of the principal moments. J Chem Phys 135, 065105 (2011).

31. Brown, J. W. et al. β-synuclein suppresses both the initiation and amplification steps of α-synuclein aggregation via competitive binding to surfaces. Scientific Reports 6, 36010 (2016).

32. Lorenzen, N. et al. The role of stable α-synuclein oligomers in the molecular events underlying amyloid formation. J Am Chem Soc 136, 3859–3868 (2014).

33. Wolff, M. et al. Quantitative thermophoretic study of disease-related protein aggregates. Sci Rep 6, 22829 (2016).

34. Pace, C. N. et al. Conformational stability and thermodynamics of folding of ribonucleases sa, sa2 and sa3. Journal of molecular biology 279, 271–286 (1998).

35. Gauhar, A., Shaykhalishahi, H., Gremer, L., Mirecka, E. A. & Hoyer, W. Impact of subunit linkages in an engineered homodimeric binding protein to α-synuclein. Protein Engineering, Design & Selection 27, 473–479 (2014).

36. Iljina, M. et al. Nanobodies raised against monomeric α-synuclein inhibit fibril formation and destabilize toxic oligomeric species. BMC biology 15, 57 (2017).

37. Waudby, C. A. et al. The interaction of alphaB-crystallin with mature alpha-synuclein amyloid fibrils inhibits their elongation. Biophys. J. 98, 843–851 (2010).

38. Månsson, C. et al. Interaction of the molecular chaperone dnajb6 with growing amyloid-beta 42 (aβ42) aggregates leads to sub-stoichiometric inhibition of amyloid formation. The Journal of biological chemistry 289, 31066–31076 (2014).

39. Kayed, R. et al. Common structure of soluble amyloid oligomers implies common mechanism of pathogenesis. Science 300, 486–489 (2003).

40. Sang, J. C. et al. Direct observation of murine prion protein replication in vitro. Journal of the American Chemical Society 140, 14789–14798 (2018).

