## Supplementary materials and data for "An engineered monomer binding-protein for *α*-synuclein efficiently inhibits the proliferation of amyloid fibrils"

### 1 Materials and Methods

**Reagents** Thioflavin T UltraPure Grade (ThT $> 95\%$) was purchased from Eurogentec Ltd (Belgium). Sodium phosphate monobasic ($NaH_2PO_4$, BioPerformance Certified $> 99.0\%$), sodium phosphate dibasic ($Na_2HPO_4$, ReagentPlus, $> 99.0\%$) and sodium azide ($NaN_3$, ReagentPlus, $> 99.5\%$) were purchased from Sigma Aldrich, UK. 1,2-Dimyristoyl-sn-glycero-3-phospho-L-serine, sodium salt (DMPS) was purchased from Avanti Polar Lipids, Inc, USA.

**Protein preparation** $\alpha$-synuclein was expressed and purified as described previously [1,2]. To determine the concentrations in solution we used the absorbance value of the protein measured at 275 nm and an extinction coefficient of 5,600 $M^{-1}$. The protein solutions were divided into aliquots, flash frozen in liquid $N_2$ and stored at -80°C, until used. AS69 was expressed and purified as previously described [3] and protein solutions were aliquoted, flash frozen in in liquid $N_2$ and stored at -80°C.

**Formation of PFF** Seed fibrils were produced at 5 mg/ml $\alpha$-synuclein in PBS buffer, as described previously [4], with the exceptions that the sonication was performed using a probe sonicator (Bandelin, Sonopuls UW 3200, Berlin, Germany) with a MS72 sonotrode, using 10% maximum power and the protein concentration was 2 mg/ml.

**Formation of seeds for elongation at pH 6.5** Seed fibrils were produced as described previously [2]. 500 $\mu$ L samples of $\alpha$-synuclein at concentrations from 500-800 $\mu$M were incubated in 20 mM phosphate buffer (pH 6.5) for 48-72 h at ca. 40°C and stirred at 1,500 rpm with a Teflon bar on an RCT Basic Heat Plate (IKA, Staufen, Germany). Fibrils were diluted to a monomer equivalent concentration of 200 $\mu$M, divided into aliquots, flash frozen in liquid $N_2$ and stored at -80°C. For experiments at pH 6.5 and $\mu$M fibril concentrations the 200 $\mu$M fibril stock was sonicated between 30 s and 1 min using a probe sonicator (Bandelin, Sonopuls HD 2070, Berlin, Germany), using 10% maximum power and a 50% cycle. For experiments at low pH with nM fibril concentrations the 200 $\mu$M stock was pre-diluted to 10 $\mu$M (monomer equivalents) in pure water, sonicated 3 times for 5 s using 10% of the maximal power and 50% cycles, using the probe sonicator.

**Formation of seeds for secondary nucleation and density gradient centrifugation** Seed fibrils were produced at the same buffer conditions, as the secondary nucleation and Density gradient centrifugation were performed. 1.2 mL sample of $\alpha$-synuclein at a concentration of 25 $\mu$M was prepared and aliquoted into 12 wells of a 96-well Half Area Black Flat Bottom Polystyrene NBS Microplate, Corning, where a single glass bead of 2.85-3.45 mm, Carl Roth, had been added. Plate was incubated at 37° for 48-72 h at 500 RPM. Sonication was performed using a probe sonicator

(Bandelin, Sonopuls UW 3200, Berlin, Germany) with a MS72 sonotrode 5 times for 1 s using 10% maximum power.

**Thioflavin T Fluorescence** Samples of 100 $\mu$ l were loaded into a 96-well Half Area Black Flat Bottom Polystyrene NBS Microplate (Corning, product number 3881). 150 $\mu$ water was added into the wells directly surrounding the wells containing sample, and the outer most wells were not used for experimental measurements. These measures minimise sample evaporation during prolonged kinetic experiments. The plate was sealed using clear sealing tape (Polyolefin Acrylate, Thermo Scientific) and placed inside a platereader (CLARIOStar or FLUOStar Omega, BMG LABTECH, Germany) that had been equilibrated to 37°C. Data points were obtained every 120-360 s, depending on the duration of the experiment. In some experiments, the fluorescence was read by averaging 12 points, measured in a ring with a diameter of 3 mm (orbital averaging mode). Excitation and emission in the CLARIOStar (monochromator) was 440 nm (15 nm bandwidth) and 485 nm (20 nm bandwidth), respectively. Excitation and emission in the FLUOStar Omega (filter) was 448 nm (10 nm bandwidth) and 482 nm (10 nm bandwidth) respectively. In addition to the proteins of interest and buffer, all samples contained 0.04 % (w/v) $NaN_3$ and 40 or 50 $\mu$M Thioflavin-T.

**Preparation of fluorescently labelled oligomers** Fluorescently labeled $\alpha$-synuclein oligomers were prepared as described previously [5,6]. In brief, we produced fluorescently labelled $\alpha$-synuclein monomer by expressing and purifying the N122C cystein variant of $\alpha$-synuclein, which was then labelled through an incubation with a 10 fold excess of Alexa 647 malimide (Thermo Fisher Scientific, Loughborough, UK), followed by removal of the excess dye with a Superdex 200 10/300 Increase gel filtration column (GE Healthcare, Amersham, UK). Wild type and fluorescently labeled N122C variant $\alpha$-synuclein were combined at a ratio of 30:1, corresponding approximately to the stoichiometry of the oligomers [7], at a total concentration of ca. 200 $\mu$M, dialysed against distilled water for 24 h and lyophilised. The dry protein was redissolved in PBS at concentrations between 500 and 800 $\mu$M and incubated at RT over night under quiescent conditions. The oligomers were then separated from the monomeric protein and larger aggregates by using a Superdex 200 10/300 Increase column that had been equilibrated with 20 mM phosphate buffer pH 7.4 and 50 mM NaCl, collecting fractions of 500 $\mu$l. The exact concentration of the oligomer fractions are difficult to determine, due to the weak absorption signal. However, based on the absorptions at 275 nm and 647 nm, we estimated the oligomer concentration to be 3-6 $\mu$M in monomer equivalents, corresponding to an oligomer number concentration of 100-200 nM, which also corresponds roughly to the concentration of Alexa label.

**AFM imaging** Atomic force microscopy images were taken with a Nanowizard II atomic force microscope (JPK, Berlin, Germany) using tapping mode in air. Solutions containing fibrils were diluted to a concentration of 1 $\mu$M (in monomer equivalents) in water and 10 $\mu$L samples of the diluted solution were deposited on freshly cleaved mica and left to dry for at least 30 min. The samples were carefully washed with 50 $\mu$L of water anid then dried again before imaging.

**Density Gradient Centrifugation** The density gradient centrifugation was performed as previously described [8].

**Thermophoresis experiments** The thermophoresis experiments with fluorescently labeled monomeric and oligomeric $\alpha$-synuclein were performed as described previously [6], using a Monolith instrument (Nanotemper, Munich, Germany) and glass capillaries (Nanotemper, Munich, Germany) with hydrophobic coating (oligomeric $\alpha$-synuclein) or uncoated (monomeric $\alpha$-synuclein). A two-fold dilution series of AS69 in 20 mM phosphate buffer pH 7.4 with 50 mM NaCl was prepared and then either 10 $\mu$l of 5x diluted oligomers (corresponding to 0.6-1.2 $\mu$M) or 1 $\mu$M labelled monomer was added to each sample of the dilution series. MST experiments were performed at 40% laser power and 75% LED power (oligomers) or 60% laser power and 20% LED power (monomers). For the calculation of the relative change in fluorescence due to thermophoresis, the cursors were set before the temperature jump followed by 5 s after the temperature jump (oligomers) and 45 s after the temperature jump (monomers).

**CD melting curves** CD melting curves were obtained as described in [9], with the exceptions that slightly higher concentrations of protein were used, and the samples were heated to 90°C rather than 80°C. The CD data was fitted directly using a two-state model in order to obtain the melting temperature, $T_m$, model as described in ( [10]):

$$y = \frac{(y_f + m_f T) + (y_u + m_u T) \cdot \exp\left(\frac{\Delta H_m}{RT} \cdot \frac{T - T_m}{T_m}\right)}{1 + \exp\left(\frac{\Delta H_m}{RT} \cdot \frac{T - T_m}{T_m}\right)} \quad (1)$$

using least-square fitting from the Python packages. `scipy.optimize.curve_fit`. $y$ is the CD signal in mdeg, $y_f + m_f T$ and $y_u + m_u T$ describes linear change in CD signal of the folded and unfolded state with respect to temperature respectively, $T$ is the temperature in Kelvin, $R$ is the ideal constant constant, and $\Delta H_m$ is the change in enthalpy at $T_m$.

**Cell culture and transfections** HEK293 cells were cultured and transfected using Metafectene as previously described [11]. A53T-$\alpha$-synuclein flexibly tagged with EGFP by the interaA53Tction of a PDZ domain with its binding motif was previously described [11,12]. WT and A53T-$\alpha$-synuclein tagged by the C-terminal and N-terminal half of venus was obtained from Prof. Tiago Outeiro (University of Goettingen, Germany).

**Immunoblots** Immunoblots were carried out 24h after transfection as previously described [11] using NP40 lysis buffer containing protease inhibitors (Pierce, Thermo Fisher Scientific) and the following primary antibodies: rabbit anti-$\alpha$-synuclein (1:500, No. 2642, Cell Signalling Technology, Danvers, USA), mouse anti-beta-tubulin (1:1000, E7, Developmental Studies Hybridoma Bank, Iowa, USA). Secondary antibodies were anti-mouse IgG (NXA931) and anti-rabbit IgG (NA934V) from GE Healthcare Life Sciences (1:10000). Among the bands around 20 kDa observed with the $\alpha$-synuclein antibody, only the upper band is considered specific and was used fA53Tor quantification (see [11] for details).

**Flow cytometry** Cells were grown in 6-well plates and used 24h after transfection. Adherent cells were washed with phosphate buffer saline (PBS) 3 times and detached with trypsin. Subsequently cells were collected in FACS tubes, centrifuged for 5 min at 2000 rpm and washed again with PBS. Cell pellets were finally resuspended in 200 $\mu$L of PBS. Flow cytometry was carried out by a FACSCalibur (BD Biosciences) using forward and sideward scatter to gate cells and a fluorescence threshold of 300 AFU to detect cells with venus (YFP) fluorescence. This threshold was determined from measurements with untransfected cells and cells expressing either the N-terminal or the C-terminal half of venus only.

**Microscopy** For classification of EGFP distribution patterns, cells were grown on coverslips and fixed 24h after transfection. The distribution of EGFP fluorescence was classified manually by a blinded observer into the categories "homogenous distribution", "containing particles" and "unhealthy" (round, condensed cells) using an Olympus IX81 fluorescence microscope (60x oil objective, NA 1.35). At least 100 cells per coverslip were classified. In each experiment, 3 coverslips were evaluated per group and the results averaged.

**Drosophila stocks** Flies expressing A53T-$\alpha$-synuclein in neurons, $w[*];;P\{w[+mC] = GAL4-elav.L\}$, $P\{w[+mC] = UAS - HsapSNCA.A53T\}$ and flies expressing GFP under control of GAL4 $w[*]; P(acman)\{w[+] = UAS - GFP\}5$ were previously described [11]. Flies expressing AS69 under control of GAL4, $w[118];;P\{w[+] = UAS - AS69\}$, were generated using standard P-element transformation (BestGene Inc). Expression of A53T-$\alpha$-synuclein fused to VN and VC in neurons was achieved by genetically crossing and recombining flies carrying GAL4 under the elav promoter and VN and VC tagged A53T-$\alpha$-synuclein under the UAS promoter. The resulting genotype of these flies is $P\{w[+mW.hs] = GawB\}elav[C155]; P[w[+] = UAS - Hsap\ SNCA[A53T] : VC]$, $PBac\{attB[+mC] = UAS - VN : Hsap\ SNCA[A53T]\}/Cyo$. Flies expressing always early RNAi, w[1118]; P$\{GD4261\}$v13673, were used as control in experiments conducted with the A53T-$\alpha$-synuclein VN/VC expressing flies. These flies have been shown to have no effect in genetic screens for modifiers in neurodegenerative disease models. Flies were raised and maintained at 25°C under a 12 hour dark/light cycle.

**Climbing assay** Virgins of the stock $w[*];;P\{w[+mC] = GAL4-elav.L\}$, $P\{w[+mC] = UAS-Hsap\,SNCA.A53T\}$ were either crossed to males $w[118];;P\{w[+] = UAS-AS69\}$, or $w[*];P(acman)]\{w[+] = UAS-GFP\}5$ (control). In the F1-progeny we selected for males with pan neural [A53T]$\alpha$-synuclein and either AS69 or GFP concomitant expression. Climbing analysis was performed 5, 15 and 25 days post eclosion as previously described [11]. For each time point and per genotype 10 flies were analyzed in 10 tapping experiments with 60 s resting interval and the results averaged. The crosses where repeated n=3 times.

**Fly head fluorescence and filter trap assay** Virgins of the stock $P\{w[+mW.hs] = GawB\}elav[C155]$; $P[w[+] = UAS{-}Hsap\,SNCA[A53T] : VC]$, PBac$\{attB[+mC] = UAS{-}VN : Hsap\,SNCA[A53T]\}/Cyo$ were either crossed to $w[118];;P\{w[+] = UAS-AS69\}$ or $w[1118];P\{GD4261\}v13673$ (control) males. In the F1-progeny we selected for males with pan neural [A53T]$\alpha$-synuclein and either AS69 or always early RNAi concomitant expression. 10 fly heads were homogenized in 100 $\mu$L RIPA buffer using the Speedmill P12 (Analytik Jena AG). The lysates were centrifuged at 12000 rpm for 10min at 4°C and the supernatant collected. For the filter trap assay equal protein amounts of RIPA fly head lysates (30 $\mu$g) were adjusted to equal volumes. An equal volume of Urea buffer (8 M) was subsequently added, samples were incubated rolling at 4°C for 1 h and sonicated in a water bath for 10 min. SDS and DTT were added to a final concentration of 2% and 50 mM. Using a dot blot filtration unit, the resulting solutions were filtered through a 0.2 $\mu$m nitrocellulose membrane (Whatman) previously equilibrated with 0.1% SDS in TBS and afterwards washed in

TBS-T. Membranes were further treated as an immunoblot described previously.

**Mouse experiments** C57BL6/J-Thy1-A30P-$\alpha$-synuclein transgenic mice [13] were housed in a pathogen-free animal facility according to the guidelines from the Federation of European Laboratory Animal Science Associations (FELASA) at 20-24°C with a 12 h light/dark cycle and with food and water ad libitum. Animal breeding and experimental procedures were approved by the district government of North Rhine Westphalia (84-02.04.2014.A321). 12-15 week old mice were used for the experiments. Pre-formed fibrils (PFF) were generated as described previously [4] (see detailed description above). AS69 protein was prepared as described above. On the day of injection, aliquots of PFF solution were diluted to a concentration corresponding to 1.4 mg/ml (98 $\mu$M) $\alpha$-synuclein protein, or respectively 1.4 mg/ml synuclein and 98 $\mu$M AS69. Solutions were sonified using a model 300VT ultrasonic homogenizer (Biologics, Inc., Manassas, VA) with 60 pulses at one second each and 10% power. Mice were anesthetized by ketamine (75 mg/kg) and medetomidine (1 mg/kg) i.p. and placed into a stereotaxic frame. 2.5 $\mu$l of solution (either PFF alone, PFF + AS69, or sterile PBS) was injected at bregma coordinates 1 mm anterior / 1.5 mm lateral into each of the following two sites, cortex (1.55 mm from dura) and striatum (2.5 mm from dura) at a rate of 0.2 $\mu$l/min. After each injection, the Hamilton syringe was left in place for 2 min before moving to the deeper site, respectively withdrawing slowly. After recovery, mice received buprenorphine (20 $\mu$l per 10 g of body weight) s.c. for analgesia. Following surgery, animals were monitored daily for physical condition. Animals were sacrificed under deep anesthesia 90 days after the injection. Brains were extracted, fixed in cold 4% paraformaldehyde (PFA) in PBS for 24 hours, cryoprotected in 30% sucrose in PBS, and frozen in methylbutane at -30°C. 30 $\mu$m coronal sections were obtained using a cryostat and stained for phosphorylated $\alpha$-synuclein (pSyn): Free-floating sections were incubated in 0.3% $H_2O_2$ for 30 minutes to block endogenous peroxidase activity and rinsed 3 times in Tris-buffered saline with Tween20 (TBS-T). Sections were blocked in 3% normal goat serum and incubated overnight at 4°C in primary rabbit anti-pSyn antibody (1:500, abcam 51253). The next day, sections were washed three times with TBS-T and incubated for 30 min at 21°C with a biotinylated goat anti-rabbit secondary antibody (1:200, Vector Laboratories BA-1000). After washing, sections were incubated in Avidin-Biotin Complex (1:100, Vectastain ABC-Kit Standard PK-6100, Vector Laboratories) for 30 min at 21°C, followed by 4 mg/ml 3,3' diaminobenzidine (DAB, Sigma Aldrich) with 1% (v/v) $H_2O_2$ in Tris buffer. Sections were mounted on glass slides, counterstained with Hemalaun, dehydrated to Xylene and coverslipped with Entellan (107961, Merck). The most reliable phenotype after PFF injections are dystrophic, pSyn-positive neurites, which we termed Lewy neurites (LN) due to the similarities with Lewy neurites observed in humans. pSyn positive structures were also observed in neuronal somata, but more difficult to discriminate from background staining. The density of LN in the striatum was quantified by a blinded investigator in the section where the anterior commissure crosses from one hemisphere to the other (corresponding to Bregma 0.14 mm in the Paxinos and Franklin 2004 mouse brain atlas) using a 63x oil objective (Zeiss), Stereo Investigator Software (MicroBrightfield Bioscience, Williston, VT), the optical fractionator method, 100 $\mu$m  100 $\mu$m counting frames and a randomly placed 200 $\mu$m  200 $\mu$m grid.

### 2 Analysis of aggregation kinetics

**Strongly seeded aggregation data at neutral pH** In the case of aggregation experiments at high concentrations ($\mu$M) of pre-formed seeds under quiescent conditions, primary nucleation and fragmentation of $\alpha$-synuclein amyloid fibrils can be neglected [2]. The aggregation kinetics were analysed as previously reported by fitting a linear function to the early times of the kinetic traces [2], with the exception that fitting was only performed after the initial decrease in fluorescence intensity, which is due to the temperature dependence of ThT fluorescence and a consequence of the thermal equilibration of the multiwell-plate prepared at room temperature. The fit was performed through 5 time points starting from the point of minimal fluorescence intensity (see supplementary figure 1). The temperature-induced decrease in fluorescence intensity is superimposed to the increase in fluorescence due to fibril elongation. Therefore, using the initial growth rates likely leads to a small but systematic underestimation of the elongation rates. This fitting procedure was performed to obtain the values of $2k_+P(0)m(0)$, where $k_+$ is the fibril elongation rate constant, $m(0)$ the initial monomer concentration and $P(0)$ the initial number concentration of fibrils. For the comparison of the rates at different concentrations of AS69, we then calculate the ratios $r$:

$$r = \frac{\left(\frac{dM(t)}{dt}\right)_{AS69}\Big|_{t\approx 0}}{\left(\frac{dM(t)}{dt}\right)\Big|_{t\approx 0}} = \frac{k_+P(0)m(0,[AS69])}{k_+P(0)m(0)} \tag{2}$$

$r$ is the ratio of the initial gradient fitted to the kinetic trace for monomer elongating fibrils in the presence of AS69 and the initial gradient fitted to the kinetic trace for monomer elongating fibrils in the absence of AS69. $P(0)$ is the initial number concentration of fibrils, which is constant, as the same stock solution of seeds was used, and $m(0)$ is the initial monomer concentrations. For the prediction in Figure 4 of the main manuscript, we calculated the equilibrium concentrations of unbound $\alpha$-synuclein, $m(0, [AS69]) = [m]_{\text{free}}$ as:

$$[m]_{\text{free}} = \frac{-([AS69]_{\text{tot}} + K_D - [m]_{\text{tot}}) + \sqrt{([AS69]_{\text{tot}} + K_D - [m]_{\text{tot}})^2 + 4K_D[m]_{\text{tot}}}}{2} \tag{3}$$

where the values obtained at different $[AS69]_{\text{tot}}$ where then used for $m(0, [AS69])$ in Equation 2. This procedure corresponds to the assumption that the only effect of the AS69 is to sequester soluble $\alpha$-synuclein. Seeded aggregation experiments at very low monomer concentrations (0.75 $\mu$M seeds) were performed in order to test whether a concentration could be determined at which no net elongation is observed (supplementary figure 2). The concentration of free monomer at which the rates of fibril elongation and dissociation are equal corresponds to the equilibrium concentration [14]:

$$k_+[m]_{\text{eq}}[P] = k_-[P] \tag{4}$$

where $k_+$ is the elongation rate constant and $k_-$ is the dissociation rate constant. The equilibrium constant of monomer addition to fibril ends therefore corresponds to the inverse of the monomer concentration at equilibrium:

$$K_{\text{eq}} = \frac{k_{-}[P]}{k_{+}[m]_{\text{eq}}[P]} = \frac{1}{[m]_{\text{eq}}} \tag{5}$$

The results of these experiments are shown in supplementary figure 2. We find that even at a concentration as low as 0.5 $\mu$M, the slight increase over time of Thioflavin-T fluorescence suggests that the fibril mass increases. This result is significant, given that the ThT fluorescence in a sample that contains only fibrils decreases over time. The fact that all samples, including that measured in the absence of added $\alpha$-synuclein monomer, show an increase in ThT fluorescence during the first hour could be explained through sedimentation processes. We have shown previously that the sedimentation of fibrils can lead to an increase in detected ThT signal if the fluorescence is read from the bottom of the multiwell plate [2]. However, the subsequent increase in fluorescence intensity over several hours at concentrations of 0.5 $\mu$M or higher suggests an increase in fibril mass, and hence that the critical concentration under these conditions is lower than 0.5 $\mu$M.

**Analysis of weakly seeded aggregation data at mildly acidic pH** Aggregation experiments were also performed at very low (nM) seed concentrations at mildly acidic pH and under quiescent conditions, where it has been shown that autocatalytic secondary nucleation of $\alpha$-synuclein amyloid fibrils plays an important role [2]. In the present study, we performed these aggregation experiments in 20 mM sodium acetate buffer at pH 5.0, well below the threshold for secondary nucleation [2]. In order to quantitatively analyse the effects that AS69 and AS69fusASN exert on secondary nucleation, we started with the following equation describing the maximum aggregation rate in the presence of autocatalytic secondary nucleation [15]:

$$r_{\text{max}} = \frac{M(\infty)\kappa}{e} \qquad\qquad \kappa = \sqrt{2m(0)^{n_2}[m(0)k_+ - k_{\text{off}}]k_2} \tag{6}$$

Where $M(\infty)$ is the long time limit of the fibrillar mass concentration, $m(0)$ is the starting concentration of monomeric $\alpha$-synuclein, $n_2$ is the effective nucleus size of secondary nucleation, $k_+$ and $k_{\text{off}}$ are the rate constants of elongation and de-polymerisation respectively, and $k_2$ is the rate constant of secondary nucleation. For our analysis, we assumed the rate of de-polymerisation to be negligible and that $M(\infty)$ was not altered by the presence of AS69. Furthermore we use the upper limit of how much monomer the AS69 could possibly sequester, which is equal to the AS69 concentration. Under these assumptions, the maximum rates relative to the case where no inhibitor was present can be described as:

$$\frac{r_{\text{max},I}}{r_{\text{max},0}} = \left(1 - \frac{I}{m(0)}\right)^{\frac{n_2+1}{2}} \tag{7}$$

Where $r_{\text{max},0}$ is the maximal aggregation rate in the absence of inhibitor, $r_{\text{max},I}$ is the maximal aggregation rate at inhibitor concentration $I$ . The values of $r_{\text{max},I}$ for each kinetic trace were found by applying the gradient function from numpy and smoothing the resulting curves using a ten-point sliding average (supplementary figure 6). The maximum of the resulting curves was taken to be $r_{\text{max},I}$. For the simulations, $n_2$ was varied in order to test whether the sequestration of monomer in conjunction with a higher effective nucleus size of secondary nucleation can explain the observed strong inhibitory effect. However, even a value of $n_2$ as high as 5 was not able to explain the strong decrease in aggregation rate as a function of increasing inhibitor concentration. Therefore, we conclude that AS69 and AS69fusASN specifically inhibit secondary nucleation by interacting with species other than the monomer.

In the main manuscript, we discuss that the efficient inhibition of secondary nucleation by AS69 is likely to stem either from an interaction of AS69 alone or of the AS69:$\alpha$-synuclein complex with an oligomeric aggregation intermediate. Given the low population of nuclei/oligomers compared to monomers during the aggregation time course, as well as the high affinity of the AS69 for monomeric $\alpha$-synuclein, its affinity for such intermediate species would have to be significantly higher than that to monomers. This can be illustrated with a simple approximation. At the end of an aggregation experiment, the fibrils typically are up to several micrometers in length, corresponding to thousands of protein molecules per fibril. Therefore, the total number of 'on pathway' oligomers that has formed during the aggregation process is three to four orders of magnitude smaller than the initial monomer concentration. In order to trap a significant fraction of these intermediates in the presence of a large excess of monomer, the affinity of AS69 to these intermediates would therefore have to be at least three orders of magnitude higher than that for monomer and hence be in the picomolar regime.

The alternative explanation, the binding of the AS69:$\alpha$-synuclein complex to the aggregation intermediate, is more plausible. A clear inhibitory effect is still observed at a ratio $\alpha$-synuclein:AS69 of 100:1, which according to the estimate above corresponds to at least one order of magnitude more AS69:$\alpha$-synuclein complex than 'on pathway'-intermediate, rendering an efficient interference with the nucleation process plausible.

Therefore, we propose a model whereby rather than requiring the binding of free AS69 to an aggregation intermediate, the AS69:$\alpha$-synuclein complex is able to incorporate into a fibril precursor and efficiently prevent it from undergoing the structural rearrangement required to transform into a growth-competent amyloid fibril.

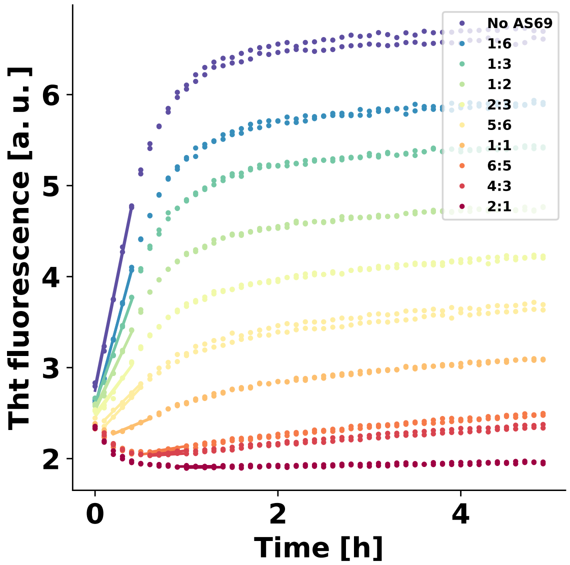


Figure 1: Linear fitting of the early times of strongly seeded aggregation kinetics. Solid lines show the fits. These data were used to produce the plot in Figure 4 c of the main manuscript. At the highest inhibitor concentrations, the rates were so low that the temperature increase upon introduction of the plate into the platereader led to an initial decrease in fluorescence intensity. Therefore, the data was fitted once the fluorescence intensity had started to increase.

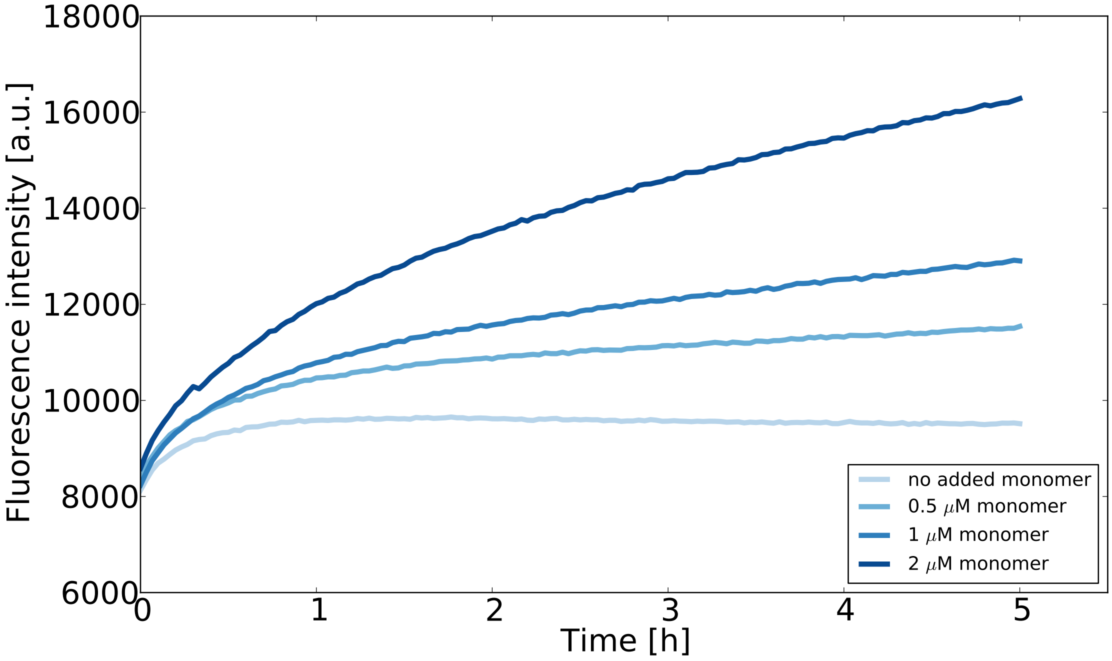


Figure 2: Seeded aggregation experiments at low monomer concentrations designed to estimate the concentration of monomeric $\alpha$-synuclein in equilibrium with fibrils. The seed concentration is in all cases 0.75 $\mu$M and the ThT concentration is 10 $\mu$M. The experiment was performed at room temperature in order to slow the reaction down and avoid temperature effects on the fluorescence upon introduction of the multiwell plate into the fluorescence platereader.

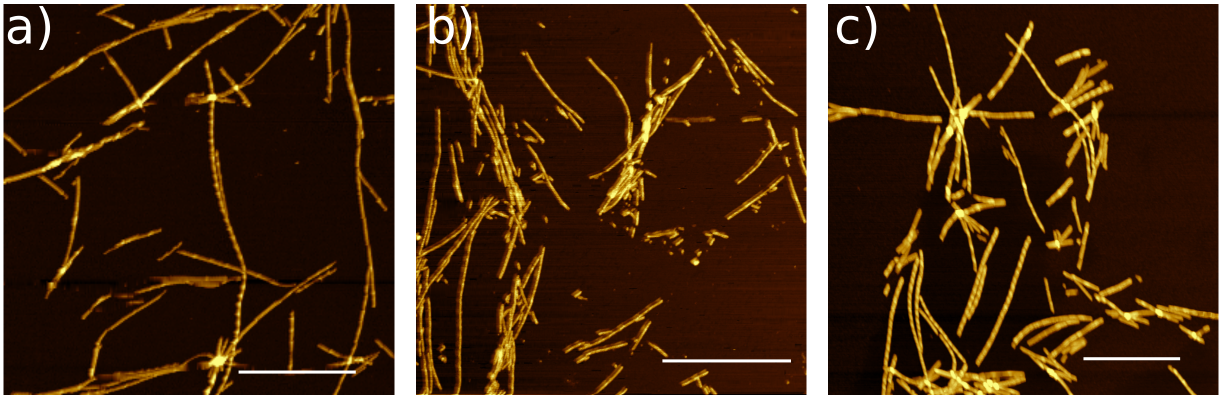


Figure 3: Characterisation of $\alpha$-synuclein fibrils formed in the presence and absence of AS69 by AFM. AFM images of 30 $\mu$M monomeric $\alpha$-synuclein that was incubated with 5 $\mu$M pre-formed fibrils, a) in the absence, b) the presence of 3 $\mu$M AS69, or c) 30 $\mu$M AS69 in 20 mM phosphate buffer at pH 6.5 under quiescent conditions at 37°C for 24 h. The scale bar represents 1 $\mu$m.

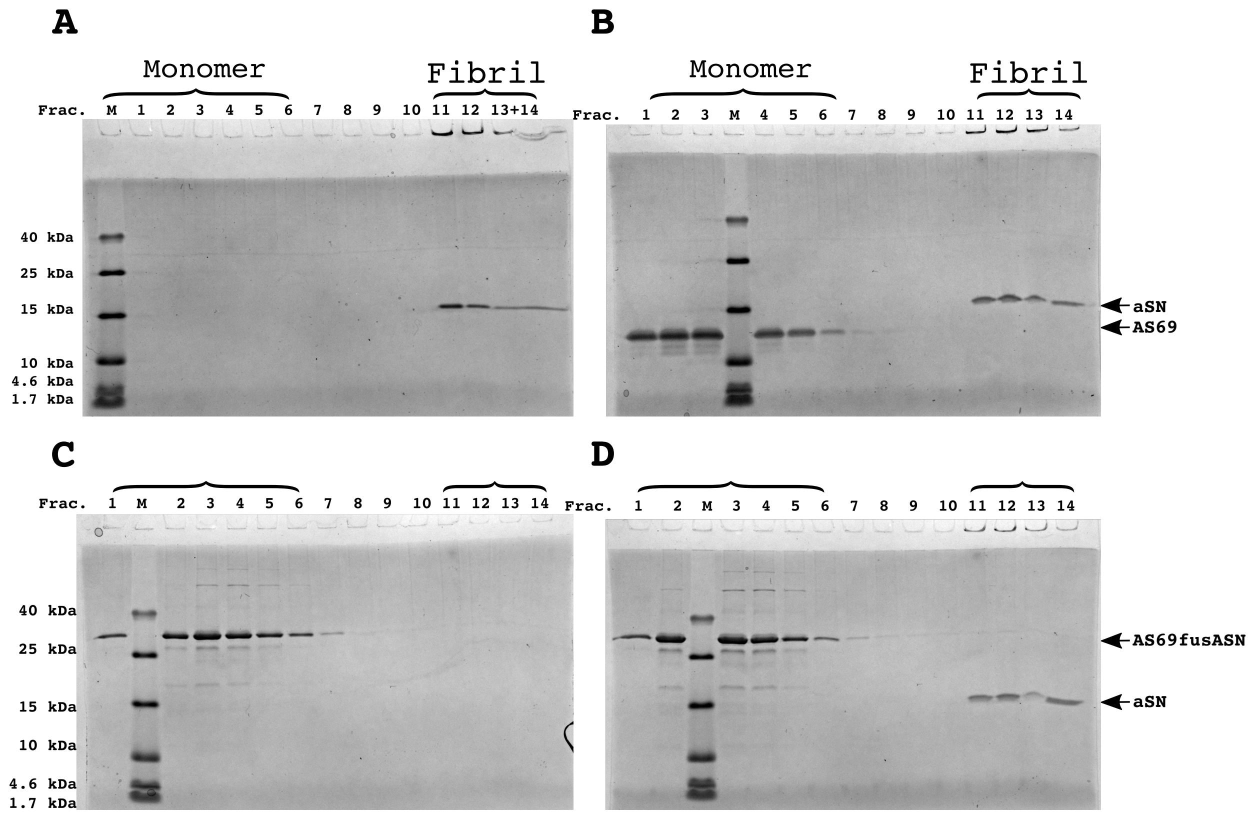


Figure 4: SDS-PAGE of Density Gradient Centrifugation (DGC) experiments to probe the binding of AS69 to $\alpha$-synulein fibrils at pH 7.4 after elongation experiments. **(a)** 25 $\mu$M seeds, **(b)** 25 $\mu$M AS69 and 25 $\mu$M seeds **(c)** 16.7 $\mu$M AS69fusASN, **(d)** 25 $\mu$M AS69fusASN and 25 $\mu$M seeds.

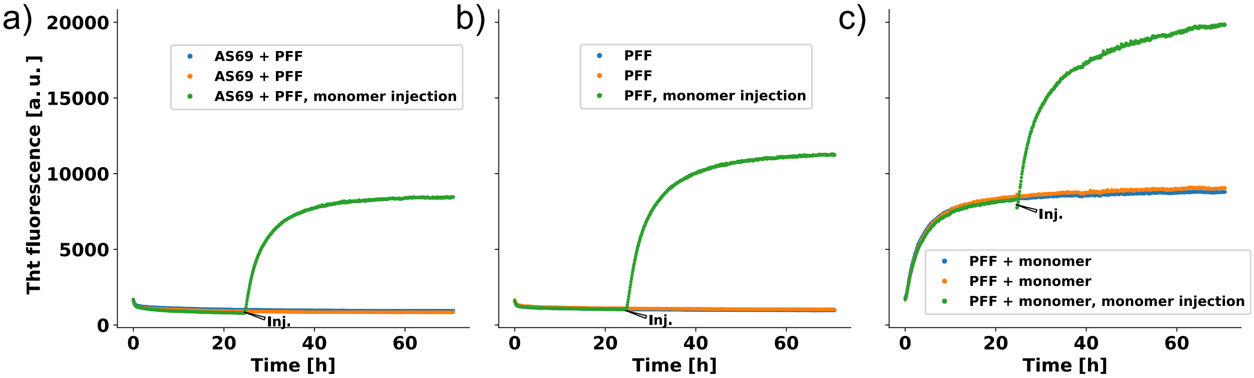


Figure 5: The effect of AS69 on the seeding efficiency of the $\alpha$-synuclein fibrils used in the mouse experiments (PFF). a) 7 $\mu$M PFF were incubated with 7 $\mu$M AS69 for 24 h and then 35 $\mu$M soluble $\alpha$-synuclein was added to the fibrils. b) 7 $\mu$M PFF were incubated for 24 h and then 35 $\mu$M soluble $\alpha$-synuclein was added to the fibrils. c) 7 $\mu$M PFF were incubated for 24 h with 35 $\mu$M soluble $\alpha$-synuclein and then an additional 35 $\mu$M soluble $\alpha$-synuclein was added to the fibrils.

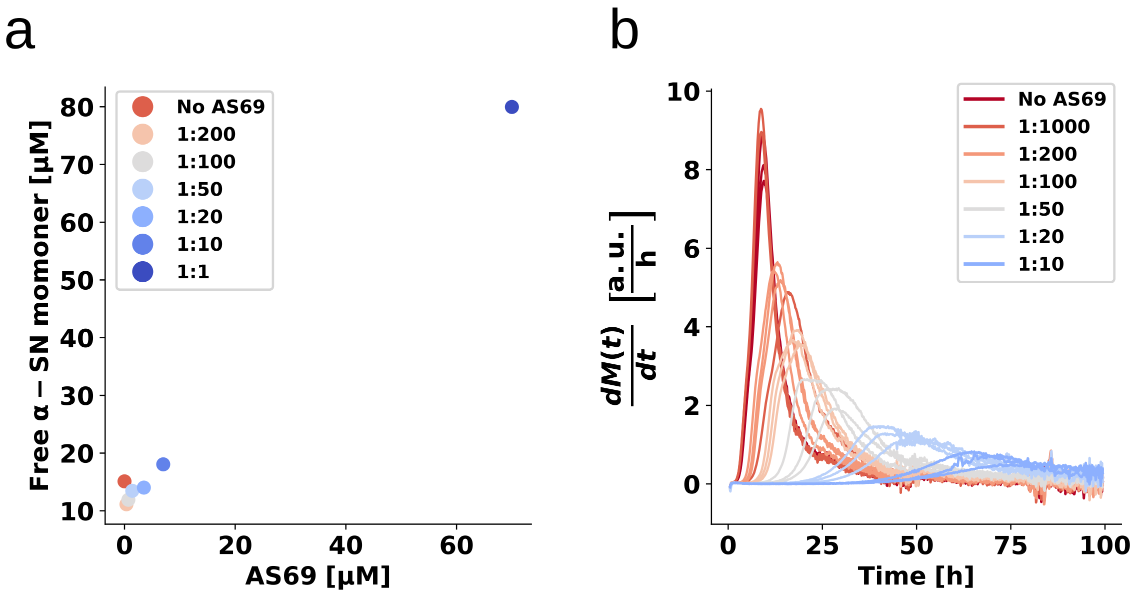


Figure 6: Weakly seeded aggregation experiments at mildly acidic pH. **(a)** The concentration of monomer left in solution after the weakly seeded experiments, as determined by the method described in [16]. **(b)** the numerically computed first derivatives (using a ten point rolling average) of the weakly seeded kinetic time courses shown in Figure 5 of the main manuscript.

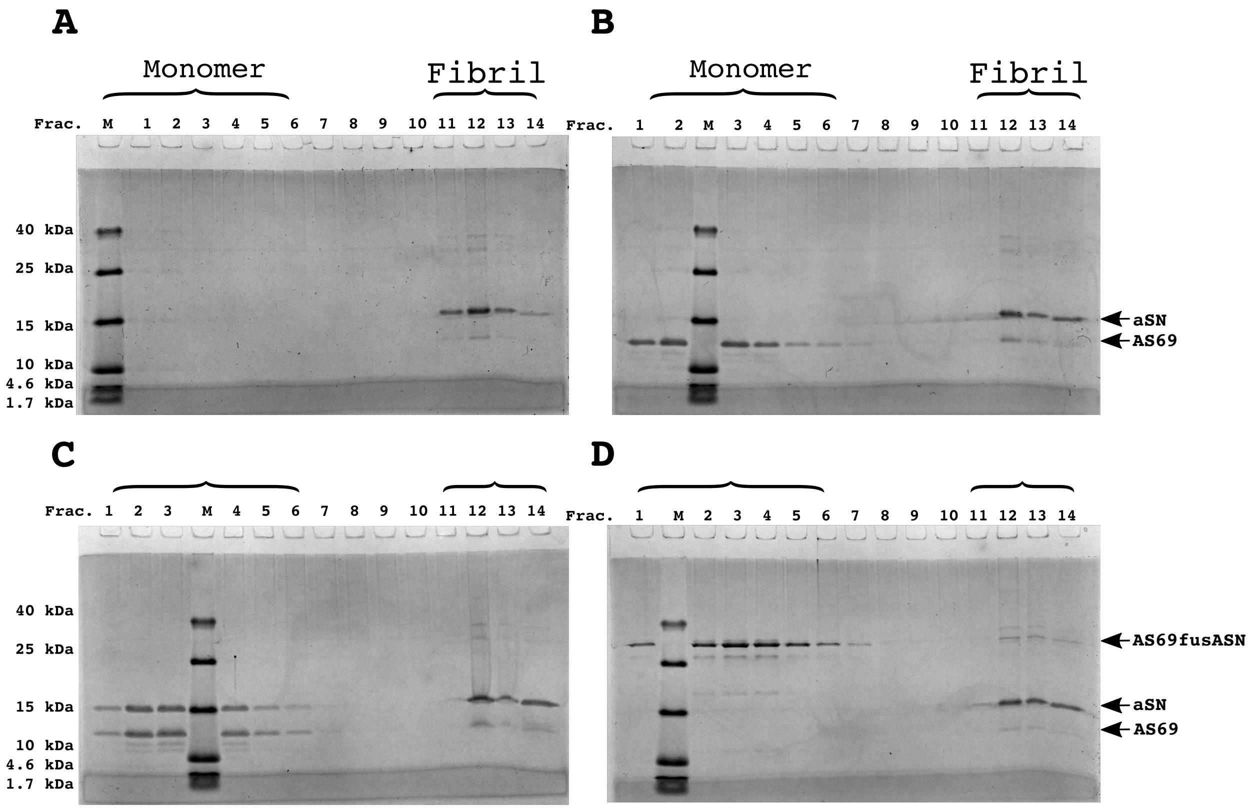


Figure 7: SDS-PAGE of Density gradient centrifugation of binding to fibril at pH 5. **(a)** 12.5 $\mu$M seeds, **(b)** 12.5 $\mu$M AS69 and 12.5 $\mu$M seeds, **(c)** 12.5 $\mu$M AS69, 12.5 $\mu$M seeds, and 12.5 $\mu$M monomer, and **(d)** 12.5 $\mu$M AS69fusASN and 12.5 $\mu$M seeds.

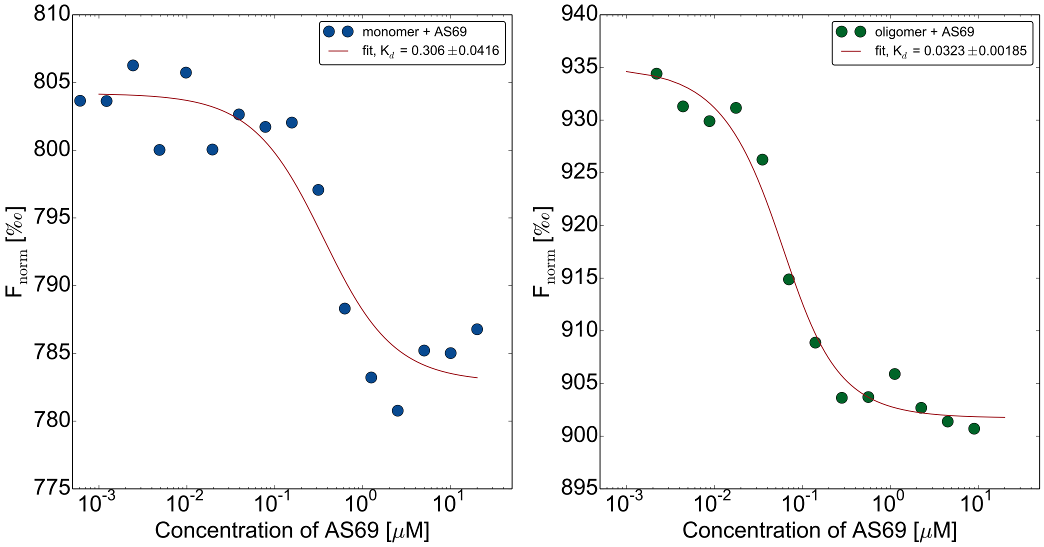


Figure 8: AS69 interacts with different $\alpha$-synuclein species. Binding of AS69 to monomeric **(a)** and oligomerc [6,7] **(b)** $\alpha$-synuclein measured by microscale thermophoresis [6].

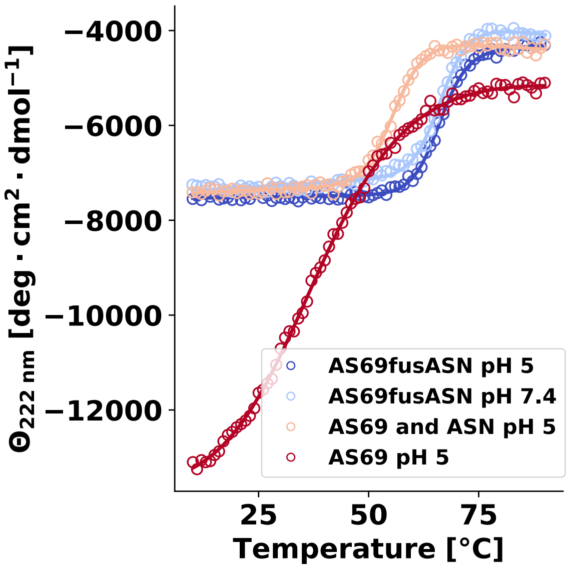


Figure 9: Melting curves of AS69 25 $\mu$M, 25 $\mu$M AS69 + 25 $\mu$M $\alpha$-synuclein in 20 mM sodium acetate pH 5.0, and 25 $\mu$M AS69fusASN 20 $\mu$M sodium acetate pH 5.0, 17.5 $\mu$M AS69fusASN 20 mM phosphate buffer 50 mM NaCl pH 7.4. Solid lines represents fit to the equation described in [10].

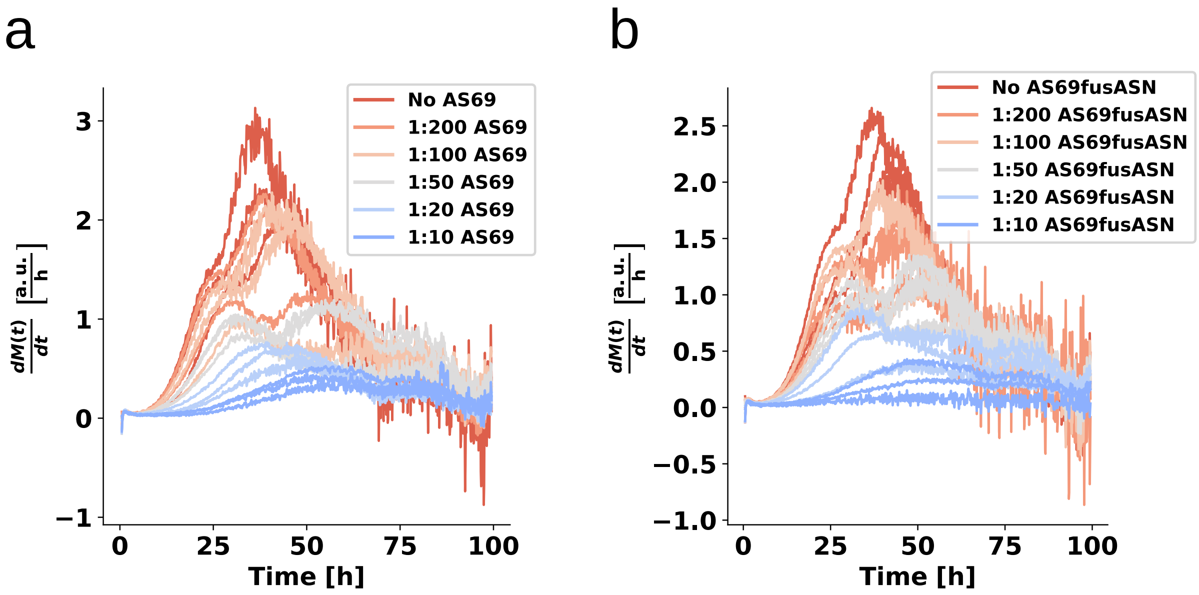


Figure 10: Weakly seeded aggregation experiments at mildly acidic pH 5. **(a)** the numerically computed first derivatives (using a ten point rolling average) of the weakly seeded kinetic time courses shown in Figure 6 c of the main manuscript. **(b)** the numerically computed first derivatives (using a ten point rolling average) of the weakly seeded kinetic time courses shown in Figure 6 **(d)**.

### References


1. Hoyer, W. *et al.* Dependence of alpha-synuclein aggregate morphology on solution conditions. *J Mol Biol* **322**, 383–393 (2002).

2. Buell, A. K. *et al.* Solution conditions determine the relative importance of nucleation and growth processes in $\alpha$-synuclein aggregation. *Proc. Natl. Acad. Sci. U.S.A.* **111**, 7671–7676 (2014).

3. Mirecka, E. A. *et al.* Sequestration of a $\beta$-hairpin for control of $\alpha$-synuclein aggregation. *Angew. Chem. Int. Ed. Engl.* **53**, 4227–4230 (2014).

4. Volpicelli-Daley, L. A., Luk, K. C. & Lee, V. M.-Y. Addition of exogenous $\alpha$-synuclein preformed fibrils to primary neuronal cultures to seed recruitment of endogenous $\alpha$-synuclein to lewy body and lewy neurite-like aggregates. *Nature protocols* **9**, 2135–2146 (2014).

5. Pinotsi, D. *et al.* Direct observation of heterogeneous amyloid fibril growth kinetics via two-color super-resolution microscopy. *Nano Lett* **14**, 339–345 (2014).

6. Wolff, M. *et al.* Quantitative thermophoretic study of disease-related protein aggregates. *Sci Rep* **6**, 22829 (2016).

7. Lorenzen, N. *et al.* The role of stable $\alpha$-synuclein oligomers in the molecular events underlying amyloid formation. *J Am Chem Soc* **136**, 3859–3868 (2014).

8. Rösener, N. S. *et al.* A d-enantiomeric peptide interferes with heteroassociation of amyloid-$\beta$ oligomers and prion protein. *The Journal of biological chemistry* **293**, 15748–15764 (2018).

9. Gauhar, A., Shaykhalishahi, H., Gremer, L., Mirecka, E. A. & Hoyer, W. Impact of subunit linkages in an engineered homodimeric binding protein to $\alpha$-synuclein. *Protein Engineering, Design & Selection* **27**, 473–479 (2014).

10. Pace, C. N. *et al.* Conformational stability and thermodynamics of folding of ribonucleases sa, sa2 and sa3. *Journal of molecular biology* **279**, 271–286 (1998).

11. Dinter, E. *et al.* Rab7 induces clearance of $\alpha$-synuclein aggregates. *Journal of neurochemistry* **138**, 758–774 (2016).

12. Opazo, F., Krenz, A., Heermann, S., Schulz, J. B. & Falkenburger, B. H. Accumulation and clearance of $\alpha$-synuclein aggregates demonstrated by time-lapse imaging. *Journal of neurochemistry* **106**, 529–540 (2008).

13. Kahle, P. J. *et al.* Subcellular localization of wild-type and parkinson's disease-associated mutant alpha-synuclein in human and transgenic mouse brain. *The Journal of neuroscience : the official journal of the Society for Neuroscience* **20**, 6365–6373 (2000).

14. Buell, A. K., Dobson, C. M. & Knowles, T. P. J. The physical chemistry of the amyloid phenomenon: thermodynamics and kinetics of filamentous protein aggregation. *Essays Biochem* **56**, 11–39 (2014).

15. Cohen, S. I. A. *et al.* Nucleated polymerization with secondary pathways. I. time evolution of the principal moments. *J Chem Phys* **135**, 065105 (2011).

16. Galvagnion, C. *et al.* Lipid vesicles trigger $\alpha$-synuclein aggregation by stimulating primary nucleation. *Nat. Chem. Biol.* **11**, 229–234 (2015).
